# Sensory tuning in neuronal movement commands

**DOI:** 10.1101/2022.11.08.515621

**Authors:** Matthias P. Baumann, Amarender R. Bogadhi, Anna F. Denninger, Ziad M. Hafed

## Abstract

Movement control is critical for successful interaction with our environment. However, movement does not occur in complete isolation of sensation, and this is particularly true of eye movements. Here we show that the neuronal eye movement commands emitted by the superior colliculus, a structure classically associated with oculomotor control, encompass a robust visual sensory representation of eye movement targets. Thus, similar saccades towards different images are associated with different saccade-related “motor” bursts. Such sensory tuning in superior colliculus saccade motor commands appeared for all image manipulations that we tested, from simple visual features to real-life object images, and it was also strongest in the most motor neurons in the deeper collicular layers. Visual-feature discrimination performance in the motor commands was also stronger than in visual responses. Comparing superior colliculus motor command feature discrimination performance to that in the primary visual cortex during steady gaze fixation revealed that collicular motor bursts possess a reliable peri-saccadic sensory representation of the peripheral saccade target’s visual appearance, exactly when retinal input is most uncertain. Consistent with this, we found that peri-saccadic perception is altered as a function of saccade target visual features. Therefore, superior colliculus neuronal movement commands likely serve a fundamentally sensory function.

## Introduction

Besides supporting a broad range of cognitive functions^1^, the superior colliculus (SC) plays a fundamental role in oculomotor control^2,3^. It issues saccade motor commands in the form of peri-movement “motor” bursts time-locked to movement onset^4-6^. Such bursts specify saccade metrics (direction and amplitude) via a distributed place code of bursting neurons^7^, and they are also widely believed to determine saccade kinematics (such as speed^8^) via a rate code within the bursts themselves. However, practically all models of saccade control by the SC rely on observations with small light spots as the saccade targets (Fig. 1A, B). Instead, in natural behavior, we generate eye movements towards image features, such as faces, cars, or trees.

**Figure 1.**
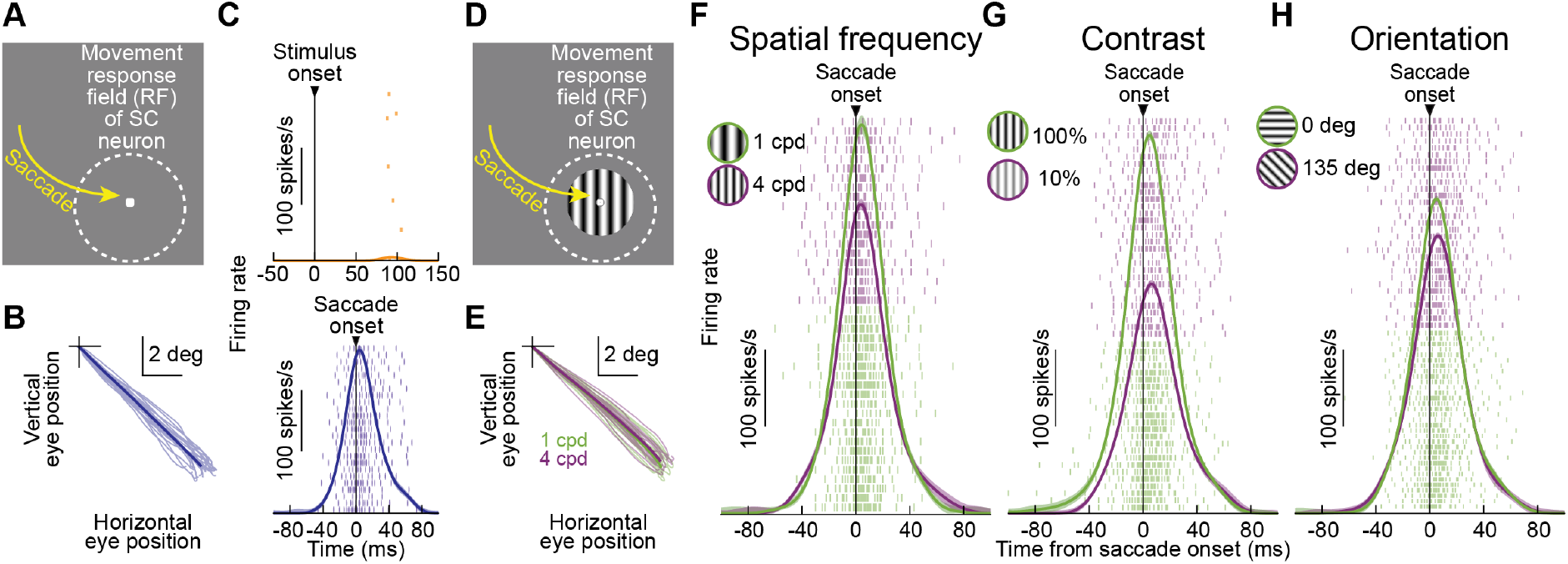
Sensory tuning in superior colliculus (SC) neuronal movement commands. **(A)** A light spot was placed within the movement-related response field (mRF) of a neuron. **(B)** Individual saccade vectors and their average (saturated line). **(C)** Stimulus-aligned (top) and saccade-aligned (bottom) firing rates from the neuron. Rasters show spike times across individual trial repetitions. The neuron exhibited practically no visual response but a strong movement-related burst. **(D)** The saccade target was now a grating. **(E)** Individual saccade vectors to a 1 cycle per deg (cpd; green) or 4 cpd grating (purple) and their averages. The saccades were matched across different images. **(F)** Nonetheless, the motor bursts of the same neuron were different for different images. **(G, H)** Similar results when we manipulated target image contrast (**G**) or orientation (**H**). Error bars: 95% confidence intervals, and numbers of trials are conveyed by the rows in the spike rasters.

If SC movement-related motor bursts were purely a neuronal control signal^8-11^, then similar saccades to different visual images should yield similar motor bursts. We tested this by training monkeys to generate saccades towards different image patches. The patches were always placed within the movement-related response field (mRF) of a recorded neuron, thus being associated with strong motor bursts. When we modified the visual features of the image patches, we also strongly modified the SC motor bursts, and for a wide range of visual image features. This modification, which also differentiated between coherent and scrambled images of real-life objects, was the outcome of a transformed representation of the peripheral saccade target visual appearance at the time of eye movement triggering. Our results document an important neuronal mechanism for bridging the trans-saccadic period of maximal sensory uncertainty caused by rapid saccade-induced retinal image shifts, and they help account for a wide range of well-known peri-saccadic alterations in visual perception.

## Results

### Saccade motor bursts are sensory-tuned

Consider the example neuron of Fig. 1C. When tested classically with a spot as the saccade target, it exhibited practically no visual sensitivity at stimulus onset (Fig. 1C, top), but it was clearly motor-related (Fig. 1C, bottom): it emitted a strong burst of action potentials time-locked to eye movement triggering. In Fig. 1D, we presented a grating inside the same neuron’s mRF, and we measured saccade-related motor bursts. In one example manipulation, we randomly varied the spatial frequency of the target across trials, and in another, we altered its contrast; in a third manipulation, we varied orientation. We always ensured that within each image manipulation, the generated saccades were behaviorally matched across the different images serving as the saccade targets (Fig. 1E; an example from the spatial frequency manipulation is shown; Methods). In every case, the neuron’s movement**-**related motor bursts were different for different images (Fig. 1F-H). This occurred even though the neuron itself was not particularly visually-sensitive (Fig. 1C), and also despite the vector-matched saccades (Fig. 1E). Thus, this example neuron’s motor bursts contained information about the visual appearance of the saccade target.

Similar observations held across our entire neuronal population, and for all image manipulations that we tested (including contrasts, spatial frequencies, orientations, and bright-versus-dark luminance contrast polarities; also see Fig. 4 below for real-life visual object images). To summarize such sensory tuning in SC neuronal movement commands, we identified, for each neuron, the image associated with the strongest (most preferred visual feature) or weakest (least preferred visual feature) “motor” burst. In every image manipulation, the difference in motor bursts between most and least preferred image features (expectedly always present by the definition of this analysis) was large, despite the matched saccades (Fig. 2A). In Fig. 2B, we also plotted raw peri-saccadic neuronal firing rates in each shown image manipulation. Even though we know that SC motor bursts can be dissociated from movement kinematics^12,13^, we also confirmed that this difference in SC motor bursts was not trivially explained by equally large differences in eye movement kinematics. We did so by plotting saccade peak velocity (Fig. 2C) from the very same trials as in Fig. 2A, B: the impact of most and least preferred trial classification on the firing rates was much larger than that on movement kinematics. We also calculated a neuronal and a kinematic modulation index for each neuron and its recorded saccades; the index was zero if there were no differences between most and least preferred trials (Methods). Neuronally, the modulation index was strongly skewed away from zero (Fig. 2D), but it straddled zero kinematically (Fig. 2E). Figure 2F also plots the modulation indices against each other, showing that they were not correlated (p=0.511, 0.059, 0.341, 0.064 in the spatial frequency, contrast, orientation, and luminance polarity manipulations, respectively; Pearson correlations: r(322)=-0.037, r(318)=0.106, r(305)=0.055, r(239)=-0.12). Thus, sensory tuning in SC motor bursts (Figs. 1, 2) was not explained by eye movement kinematics.

**Figure 2.**
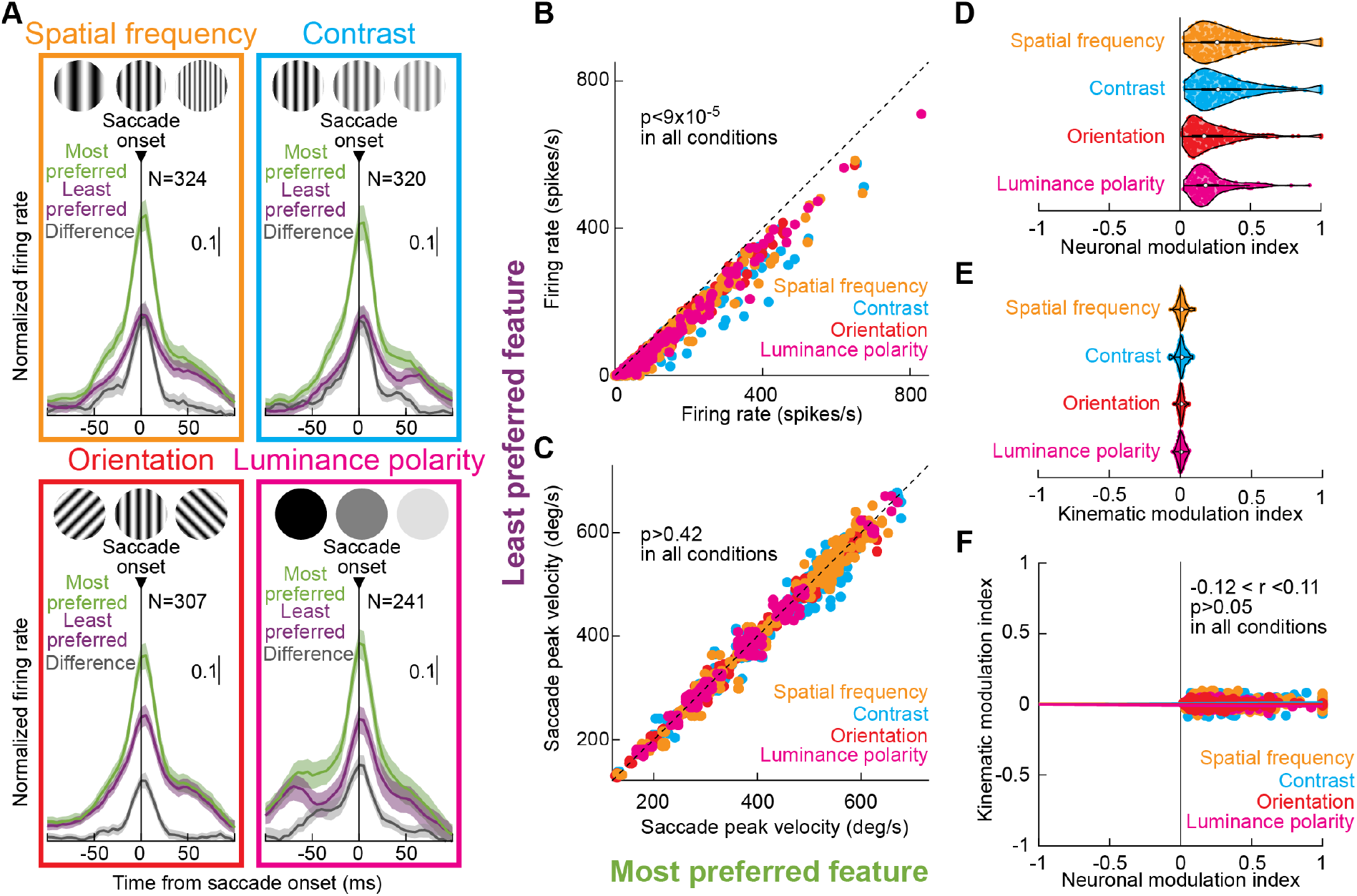
Robustness of sensory tuning in SC neuronal movement commands across image manipulations. **(A)** Average normalized population firing rates for the most (green) and least (purple) preferred stimuli within each image manipulation and their differences (gray). In all image manipulations, we observed a robust difference in SC motor bursts for different images. Note that for luminance polarity, the task was an immediate, reflexive visually-guided saccade task. Thus, visual and motor bursts occurred in close temporal proximity, explaining secondary elevation in firing rates at approximately −75 ms from saccade onset. Error bars: 95% confidence intervals, and neuron numbers are indicated in each panel. **(B)** Individual neuron raw peri-saccadic firing rates (−50 to 25 ms from movement onset; Methods) for most and least preferred targets, color-coded by each image manipulation as in **A. (C)** From the same trials as in **A, B**, saccade peak velocities were very similar for most and least preferred images, despite the large SC motor burst differences (**A, B**) (p-values indicate rank-sum test results in each image manipulation). **(D)** Violin plots of neuronal modulation indices between most and least preferred features (Methods) in the different image manipulations. Note how contrast and spatial frequency had the strongest neuronal effects and orientation had the weakest ones. **(E)** Kinematic modulation indices from the very same trials of each neuron. Despite the large neuronal modulations (**D**), the kinematic modulations straddled zero, consistent with **C. (F)** Neuronal and kinematic modulation indices were not correlated, suggesting that our results were not explained by differences in saccades for different images. Supp. Fig. 1 also shows that other behavioral properties of saccades did not explain our neuronal results. Neuron numbers in **B**-**F** are the same as in **A**. Also see Fig. 4 showing sensory tuning in SC neuronal movement commands with real-life object images.

Naturally, the SC’s population code^7^ can also alter SC motor bursts. For example, a slightly deviated saccade vector for one image could be associated with an altered neuronal response, simply by activating a different portion of a given neuron’s mRF. We minimized this by a clear marker at the center of every image (e.g. Fig. 1D; Methods), to better guide saccades. More importantly, we also performed post-hoc vector matching of the saccades before analyzing the motor bursts, removing any outlier eye movement vectors (Methods; see Fig. 1E). To confirm the effectiveness of such vector matching, we tested saccade metrics in the accepted trials. For each neuron, and for the very same trials as in Fig. 2, we plotted saccade amplitude and direction errors, as well as their differences, for the most and least preferred images. The results, shown in Supp. Fig. 1, confirm that saccade metric differences do not trivially explain the large SC motor burst differences of Figs. 1, 2.

It might additionally be argued that the intrinsic salience of individual image features might have introduced an internal reward signal^14,15^ associated with some saccade targets versus others. However, reward expectation affects both SC activity^16^ (globally, including visual bursts, delay-period activity, and motor bursts) and the eye movement properties themselves^14^, whereas we saw minimal kinematic alterations with large neuronal effects (Fig. 2, Supp. Fig. 1; also see refs. ^12,13^ for further evidence of dissociation between motor bursts and kinematics in other contexts). More importantly, we equalized the image conditions as much as possible by enforcing a time delay between stimulus onset and the instruction to make a saccade (Methods). Besides ensuring a stable scene image at the time of saccade triggering, which is more similar to natural looking behavior, this enforced delay reduced the impacts of potential differences in intrinsic image salience. For example, saccadic reaction time distributions were strongly overlapping for different spatial frequencies (Supp. Fig. 2A), unlike what would happen with no enforced fixation^17,18^; similar results were also obtained in our other image manipulations (Supp. Fig. 2B, C), suggesting that potential intrinsic image salience did not explain the results of Figs. 1, 2. Indeed, in one image manipulation (luminance polarity), we also explicitly skipped enforced fixation (Methods). Both example neurons (Supp. Fig. 3A, B) and the population (Fig. 2, luminance polarity data) showed the same sensory tuning in SC neuronal movement commands even though saccadic reaction times were different across different conditions in this data set (ref. ^19^ and Supp. Fig. 3C). Therefore, whether stimuli were intrinsically salient (e.g. having low spatial frequency or high contrast) or not, sensory tuning in SC neuronal movement commands was still present.

**Figure 3.**
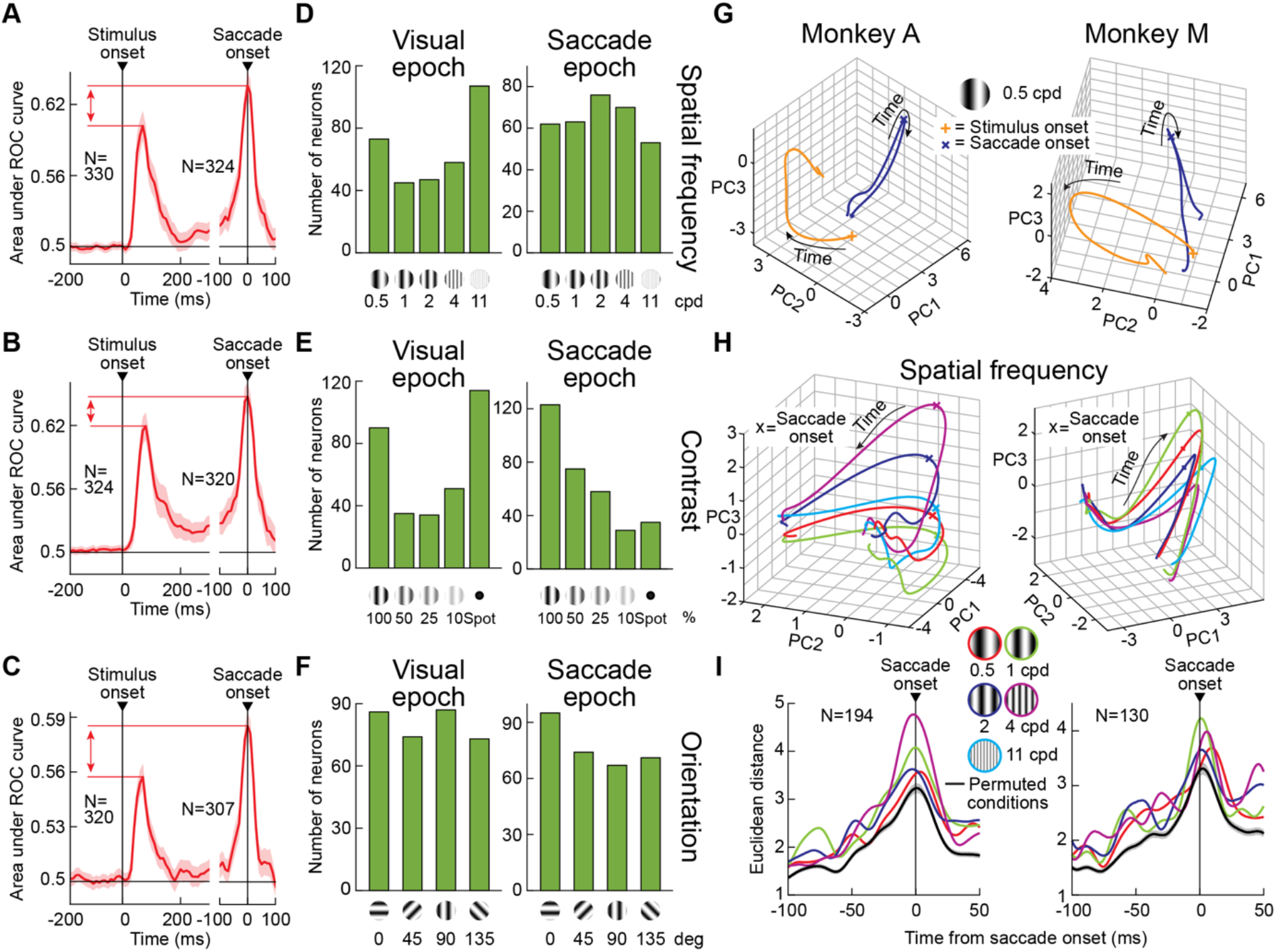
Transformation of the visual sensory representation of saccade targets in the SC at the time of eye movement generation. **(A-C)** Area under the ROC curve (AUC) discrimination performance between most and least preferred images as a function of time from stimulus or saccade onset: (**A**) spatial frequency; (**B**) contrast; (**C**) orientation. In each case, it was possible to better discriminate between most and least preferred images from the motor bursts of individual SC neurons than from the visual responses (vertical red arrows). Error bars: 95% confidence intervals across neurons, and neuron numbers are indicated in each panel. Note that for saccade burst analyses, we sometimes excluded a small number of neurons if there were not enough vector-matched movements across all image conditions of a given manipulation (Methods). This explains the small difference in neuron numbers between visual and motor burst epochs. **(D-F)** In each image manipulation, distribution of preferred features in the initial visual response (left column; visual epoch) or in saccade motor bursts (right column; saccade epoch). Sensory tuning in SC neuronal movement commands increased representation of visual features that were not normally preferred in the visual epochs. **(G)** For an example image (0.5 cpd grating), trajectory of each monkey’s entire population of neurons’ firing rates after stimulus onset (orange; 0-200 ms) or peri-saccadically (dark blue; −100 to 50 ms), after dimensionality reduction using PCA (Methods). Peri-saccadic population activity was more spatially-constrained, consistent with recent evidence^26^. Importantly, population activity occupied almost orthogonal manifolds in the visual and motor burst epochs, suggesting a transformed representation, consistent with **A**-**F. (H)** When we plotted the motor burst epochs (with higher zoom and a different view), but now for all the different spatial frequency images, the population trajectories were differentiated for each feature. Thus, it is possible for recipient neurons to read-out information about saccade target visual appearance from the SC motor bursts. **(I)** We picked a reference peri-saccadic trajectory (11 cpd) in the high-dimensional population activity space of each monkey’s neurons, and we then plotted Euclidean distances of population activity in each other image feature from this trajectory. For each image and monkey, Euclidean distances peaked at saccade onset (consistent with Figs. 1-2), and they were higher than Euclidean distances obtained with randomly shuffled reference and non-reference trajectories (black). Thus, individual features were discriminable within SC motor bursts. Supp. Figs. 7-8 show similar population analyses from all of our other image manipulations.

### Stronger effect than in SC visual bursts

To quantitatively test discrimination performance of the most and least preferred image features from the visual or motor bursts of individual SC neurons, we used area under the ROC curve (AUC) analyses (Methods), like in visual cortical pre-saccadic analyses^20^. In Fig. 3A-C (left), we calculated AUC at every time bin relative to stimulus onset, comparing a neuron’s distribution of firing rates for most and least preferred images during steady-state gaze fixation. We defined the most and least preferred images in the visual bursts similarly to how we defined them for motor bursts: the most or least preferred image was that evoking the strongest or weakest visual response by a neuron, respectively (Methods; raw population firing rates are shown in Supp. Fig. 4, formatted similarly to Fig. 2). We only focused on the conditions enforcing a delay between stimulus onset and saccade generation, to avoid a temporal mixing between visual and motor bursts in reflexive saccade paradigms. In Fig. 3A-C (right), we performed the same analysis around saccade onset. In the visual epoch, there was an expected peak in AUC value soon after stimulus onset; SC visual responses represent visual-scene image information^17,21^. Critically, across the population, and in all image manipulations, there was also a strong peri-saccadic peak in AUC discrimination performance, consistent with Figs. 1, 2. This peak was at least as high as that in pre-saccadic elevations of visual cortical activity in area V4^20^. Thus, SC saccade motor bursts possess robust information about the visual appearance of the peripheral saccade target.

Intriguingly, we could better discriminate the most and least preferred images from SC motor rather than visual bursts (compare visual and motor epoch peaks in Fig. 3A-C; error bars denote 95% confidence intervals). These results likely represent a yet-to-be-appreciated underlying neuronal mechanism for well-known pre-saccadic enhancements of visual perception relative to steady-state gaze fixation^22-25^.

It is also important to note that a peak in peri-saccadic AUC discrimination performance also clearly emerged in the reflexive saccade version of our tasks. Specifically, in this task, individual neurons preferred stimuli of different luminance polarity and contrast in their motor bursts (Fig. 2, luminance polarity data), with similar AUC discrimination performance levels as those seen in the other image manipulations (Supp. Fig. 5A). Therefore, sensory tuning in SC neuronal movement commands still persisted for reflexive orienting responses.

### Transformed peri-saccade SC scene coding

Across all image manipulations shown so far, and irrespective of delayed or reflexive saccades, our results document a robust sensory signal embedded within SC motor bursts, which is not fully accounted for by saccade metric and kinematic properties, and which is at least as good as that present in initial visual sensory responses (and also in pre-saccadic visual cortical activity^20^). We next investigated which image features individual neurons preferred.

In the visual burst epochs, we observed expected SC sensory tuning properties. For example, SC neuron visual bursts expectedly^17,27-29^ preferred low spatial frequencies (Fig. 3D) and high contrasts (Fig. 3E), with the caveat being that the spot at each patch center (used to improve saccade vector matching; Methods) interacted with the underlying image patch, especially when the patch was least visible (e.g. having low contrast or high spatial frequency). The spot (introducing a broadband spatial frequency signal), therefore, changed the spectral and luminance content of the overall image, and this increased the numbers of neurons preferring high spatial frequencies or low contrasts in Fig. 3D, E when compared to the literature ^17,27-29^. This, in itself, further confirms that our neurons behaved as expected in their stimulus-evoked visual sensory responses.

Peri-saccadically, all image features were well represented by the motor bursts (Fig. 3D-F, saccade epoch). Critically, neurons often changed their preferred image features relative to the initial visual burst epoch (see example neurons in Supp. Fig. 6). The result, across the population, was a transformed visual representation in the motor bursts, evidenced by different distributions of preferred image features across neurons (compare visual and saccade epochs in Fig. 3D-F). Importantly, unlike in visual responses, mid-spatial frequencies became more prevalently represented in the motor bursts (Fig. 3D). Similarly, for the contrast manipulation, lower contrasts (e.g. 25%) became preferred more often (Fig. 3E). Thus, there was effectively some amount of equalization of preferred features in the motor bursts: images that were less likely to be preferred in visual bursts were more preferred during the motor commands.

The above transformation of visual information in the SC motor bursts is very interesting given that spatial frequency perception in humans increases its bandwidth (shifting towards higher spatial frequencies; like in Fig. 3D) pre-saccadically^25^. Pre-saccadic contrast sensitivity is also enhanced^24,28^, consistent with the enhanced preferences for low contrasts in our SC motor bursts. In fact, in our reflexive saccade paradigm, we used a small saccade target (as opposed to large gratings; Methods). Even in that case, a clear preference for high contrasts in the visual response epochs was transformed into one in which lower contrasts became much better represented by the SC population at saccade triggering (Supp. Fig. 5B, C; also see the example neuron of Supp. Fig. 3B). Therefore, the visual appearance of the saccade target is represented by SC motor bursts using a transformed code amplifying weak visual signals.

To further understand the transformation of visual target representations in SC motor bursts, we estimated the high-dimensional state-space trajectory^26^ of our recorded neurons’ activities either after stimulus onset or peri-saccadically (Methods). We first created, for each time, a point in an N-dimensional space of N recorded neurons. For visualization purposes, we then performed principal components analysis (PCA) and plotted the 3-dimensional trajectory of population activity using the first 3 principal components, which accounted for a great majority of the variance in the neuronal data (Methods). Prior work suggested that the population trajectory in the motor burst epoch should be relatively straight, compared to that in the visual burst epoch, suggesting a temporal alignment of the population at the time of saccade triggering^26^. We confirmed this (Fig. 3G). Critically, however, we uncovered two additional key properties of the SC population activity that are particularly relevant here.

First, there was an almost orthogonal relationship between population trajectory during the visual and motor burst epochs (Fig. 3G). Thus, areas reading out SC population activity encounter largely different state-space loci during fixation and saccades, consistent with the altered feature preferences of Fig. 3D-F.

More importantly, when we repeated the same PCA analyses around saccade onset, but now differentiating trials based on the different presented images, we confirmed the results of Figs. 1, 2, 3A-F: peri-saccadic population trajectory was different for different saccade target images, despite the matched vectors across all spatial frequency conditions (Fig. 3H). To demonstrate that feature information was indeed present in the population during motor bursts, we then used one image feature (11 cpd in the example of Fig. 3I) as a reference high-dimensional population trajectory. Then, we calculated the Euclidean distance, at every time point, between each other feature condition (e.g. 4 cpd) and this reference condition. The Euclidean distance always peaked at motor burst time, and it was different for different image features (Fig. 3I). Moreover, all of the peak Euclidean distances were larger than those expected by random permutation of reference and condition trajectories (black line in Fig. 3I; Methods). Supp. Fig. 7 shows how Euclidean distances were consistently different for different images in our other image manipulations as well. Therefore, at the time of saccades, there was a graded, differentiated representation of visual image features by SC populations.

### Visual objects are also well-represented

Since natural looking behavior typically involves foveating specific objects, a strong test of the ecological relevance of sensory tuning in SC neuronal movement commands would be to check high-level visual object representations. We, therefore, studied motor bursts for luminance-equalized natural images of animate and inanimate objects, as well as feature- and spectral-scrambled versions of them (Fig 4A; Methods). We recently found that SC visual responses are indeed sensitive to coherent visual object images^30^, so we wondered whether this also held in the motor bursts.

**Figure 4.**
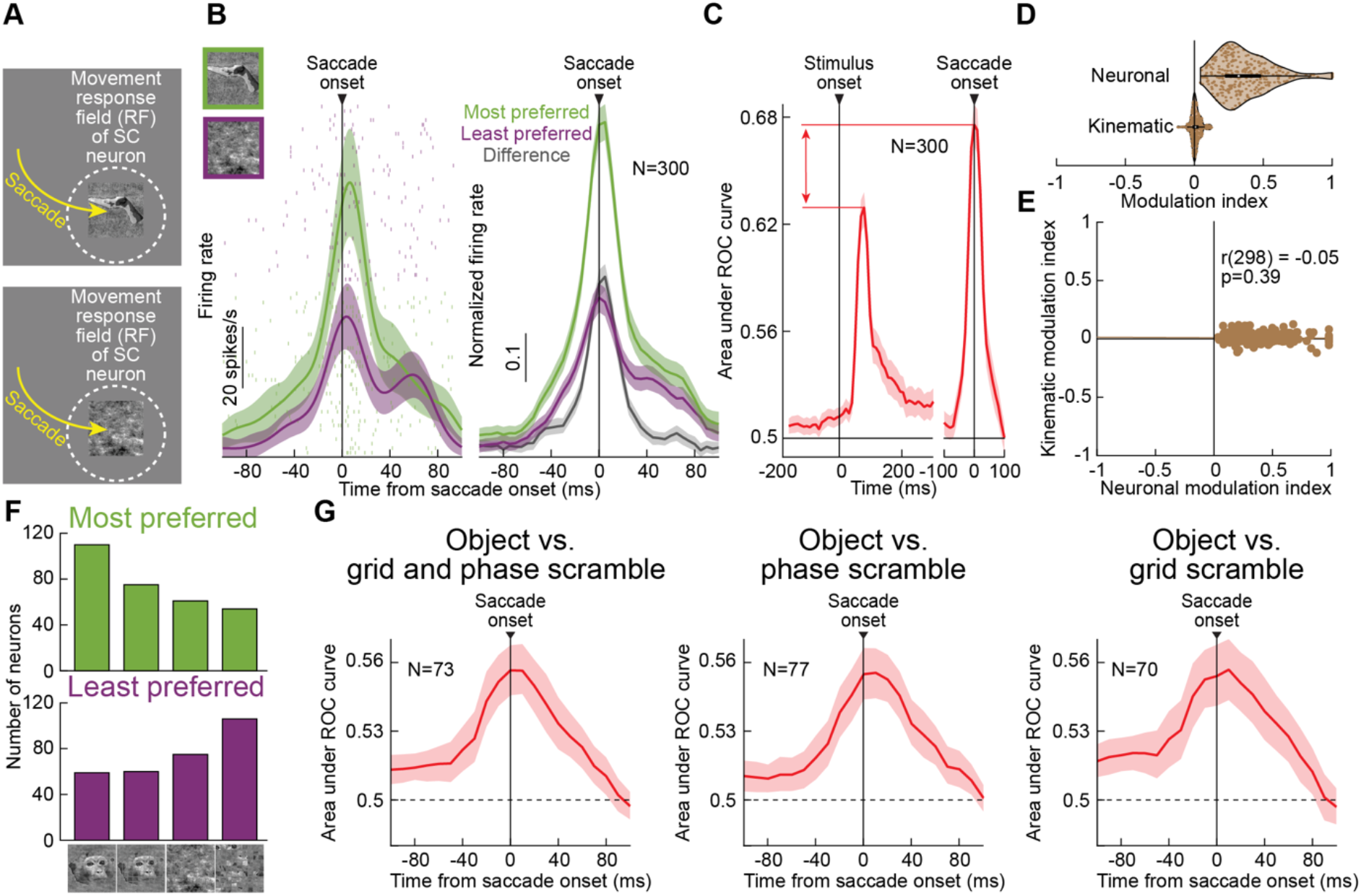
Sensitivity of SC neuronal movement commands to coherent visual object images. **(A)** We tested saccades to real-life objects or scrambled versions of them^30^. **(B)** Left: example neuron’s saccade motor bursts for an image of a hand (green) or a phase-scrambled^30^ version of it (purple); the neuron preferred the coherent object image. Right: across the population, most preferred images (green) had a much higher peri-saccadic firing rate than least preferred images (purple), and the difference (gray) was larger than in our other image manipulations (compare to Fig. 2A). **(C)** Like in Fig. 3A-C, AUC discrimination performance was higher in motor bursts than in visual bursts. **(D, E)** Moreover, there was no correlation between neuronal and kinematic modulation indices across the population (as in Fig. 2D-F for simpler image features). Supp. Fig. 8A-C shows additional controls for other saccade behavioral metrics. **(F)** In the SC motor bursts, the most preferred images were most likely to be coherent object images (first two columns; top histogram), and the least preferred images were most likely to be scrambles (last two columns; bottom histogram). Also see Supp. Fig. 8D for further evidence from high-dimensional population state-space analyses. **(G)** Consistent with this, we found a substantial number of neurons with significant (Methods) AUC discrimination performance between real and scrambled images of different kinds in the saccade motor bursts. Thus, SC motor bursts are sensitive for real-life object images as saccade targets. Error bars in all panels: 95% confidence intervals.

The largest differences in SC motor burst strengths between most and least preferred images occurred with such naturalistic images. For example, Fig. 4B shows that the motor burst of an example neuron was almost completely abolished when making saccades towards a non-preferred scrambled image. Across the population, there was a larger difference between preferred and least preferred images in this experiment than with simple feature dimensions (compare Fig. 4B, right to Fig. 2A), and AUC discrimination performance was also still higher during the motor bursts than during earlier visual bursts (Fig. 4C). These effects were, again, not explained by eye movement effects (Fig. 4D, E; Supp. Fig. 8A, B).

Most interestingly, a majority of neurons had the most preferred image during saccade motor bursts as a real object image (first two columns in the histogram of Fig. 4F, top) and the least preferred image as a scramble (last two columns in the histogram of Fig. 4F, bottom). Individual SC neurons’ motor bursts also contained significant information about whether a saccade target was a coherent or scrambled object image, as revealed by a significant peak in peri-saccadic AUC discrimination performance in Fig. 4G. And, high-dimensional population trajectories in the motor burst epoch differentiated between coherent and scrambled object images (Supp. Fig. 8D).

Thus, sensory tuning in SC neuronal movement commands extended to high-level visual object representations. This suggests an ecological relevance of sensory tuning in SC neuronal movement commands in more naturalistic active vision scenarios.

### Strongest effect in least visual neurons

Sensory tuning in SC neuronal movement commands was not restricted to a single functional cell type, but it was a robust property of all movement-related neurons. In fact, AUC discrimination performance was highest for the most motor neurons (occupying the deeper SC layers^5,6^). In each task (with dissociated visual and motor burst times), we repeated our AUC analyses of Fig. 3A-C, but now after classifying how each neuron responded in either the visual or motor epoch of the task (Methods). AUC discrimination performance in the motor bursts was always the highest for the most motor neurons and decreased progressively for visual-motor and then visual neurons (Fig. 5B-E; note that Fig. 5A shows example normalized population firing rates clarifying the different functional cell types). More motor neurons also showed sensory tuning in our reflexive saccade paradigm with small targets (see the example neuron of Supp. Fig. 3A).

**Figure 5.**
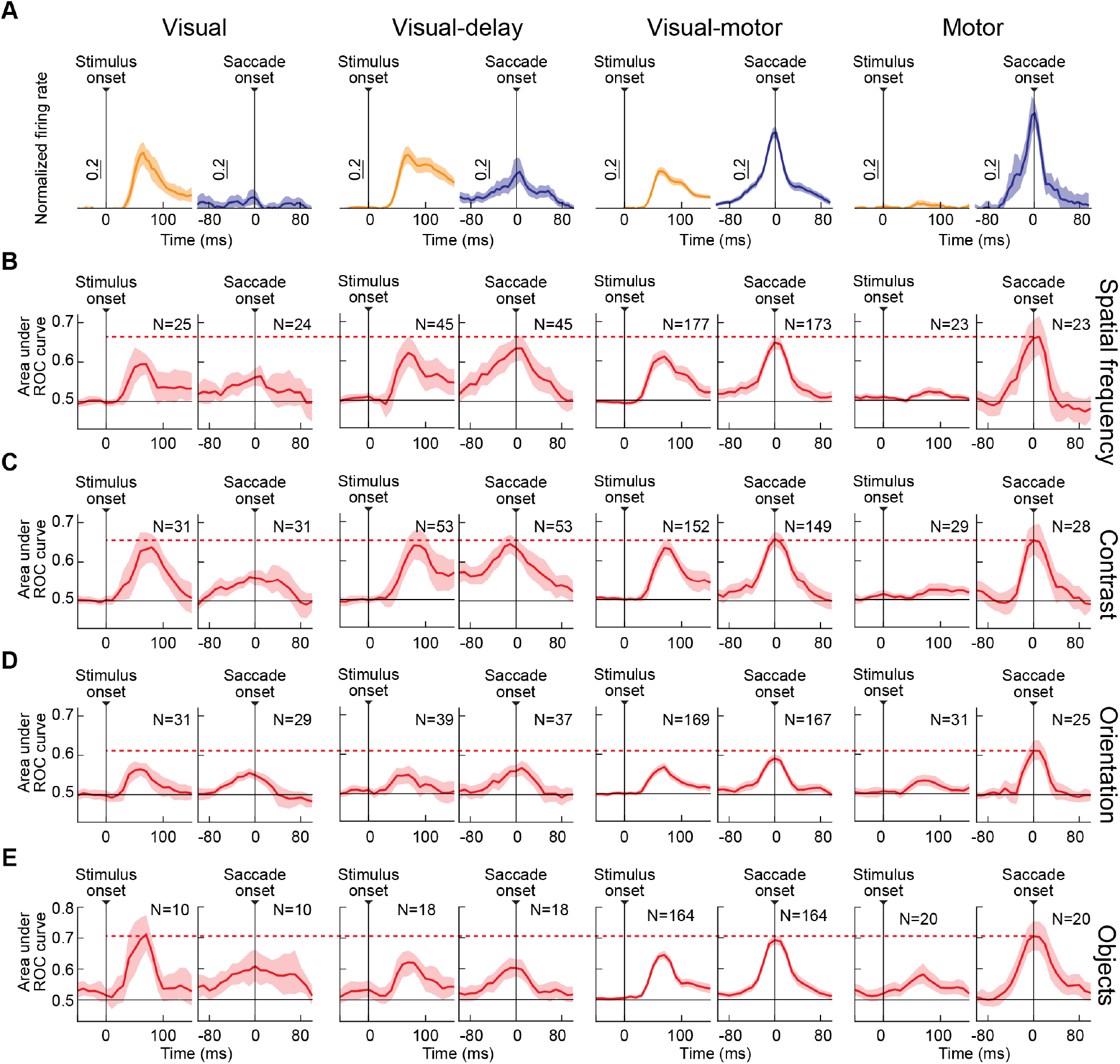
Pervasiveness of sensory tuning in SC neuronal movement commands across movement-related cell types. **(A)** In each image manipulation, we classified neurons as visual (leftmost column), visual-delay (second column), visual-motor (third column), or motor (fourth column) based on their peri-stimulus, peri-saccadic, or delay-period activity (Methods). The example shown is for the preferred conditions of the spatial frequency manipulation. **(B-E)** For each image manipulation, we plotted peri-stimulus or peri-saccadic AUC discrimination performance (like we did in Figs. 3, 4) but for each functionally-identified cell type separately. Visual neurons had the weakest peri-saccadic discrimination performance because they had weak or non-existent motor bursts. However, for all other cell types, discrimination performance in the motor bursts was at least as good as, if not better, than discrimination performance in the visual bursts. More importantly, motor burst discrimination performance was the highest in each image manipulation for the most motor neurons (see horizontal dashed line comparing motor neuron peri-saccadic performance to the other cell types). Error bars in all panels: 95% confidence intervals.

Local field potential (LFP) analyses also confirmed that peri-saccadic LFP modulations in the deeper (more motor) SC layers were different for different images as the saccade targets, again for metrically and kinematically matched saccades (Supp. Fig. 9A-D). LFP modulations also distinguished between real objects and scrambled versions of them (Supp. Fig. 9E-G). Thus, as part of the transformation in population representation of images alluded to above (Fig. 3), an increasingly strong sensory tuning emerged in the most motor SC layers.

### Relation to V1 effect in steady fixation

Why sensory tuning in SC neuronal movement commands? In seminal work, Sommer and Wurtz^31^ reported that SC motor bursts are relayed faithfully to the cortex. Our results imply that, besides the intended saccade vector, SC-sourced corollary discharge can provide a visual preview of the soon-to-be-foveated target, allowing the visual system to bridge a gap of sensory uncertainty caused by rapid eyeball rotation. Indeed, when we simultaneously recorded V1 activity in the same tasks of Fig. 1 (Methods), we found that AUC discrimination performance in SC motor bursts peaked peri-saccadically to a level that was at least as good as how V1 neurons discriminated between their most and least preferred peripheral images during steady-state gaze fixation (Fig. 6; the orange SC curves show results from the SC neurons that were recorded simultaneously with the V1 neurons, with consistent results).

**Figure 6.**
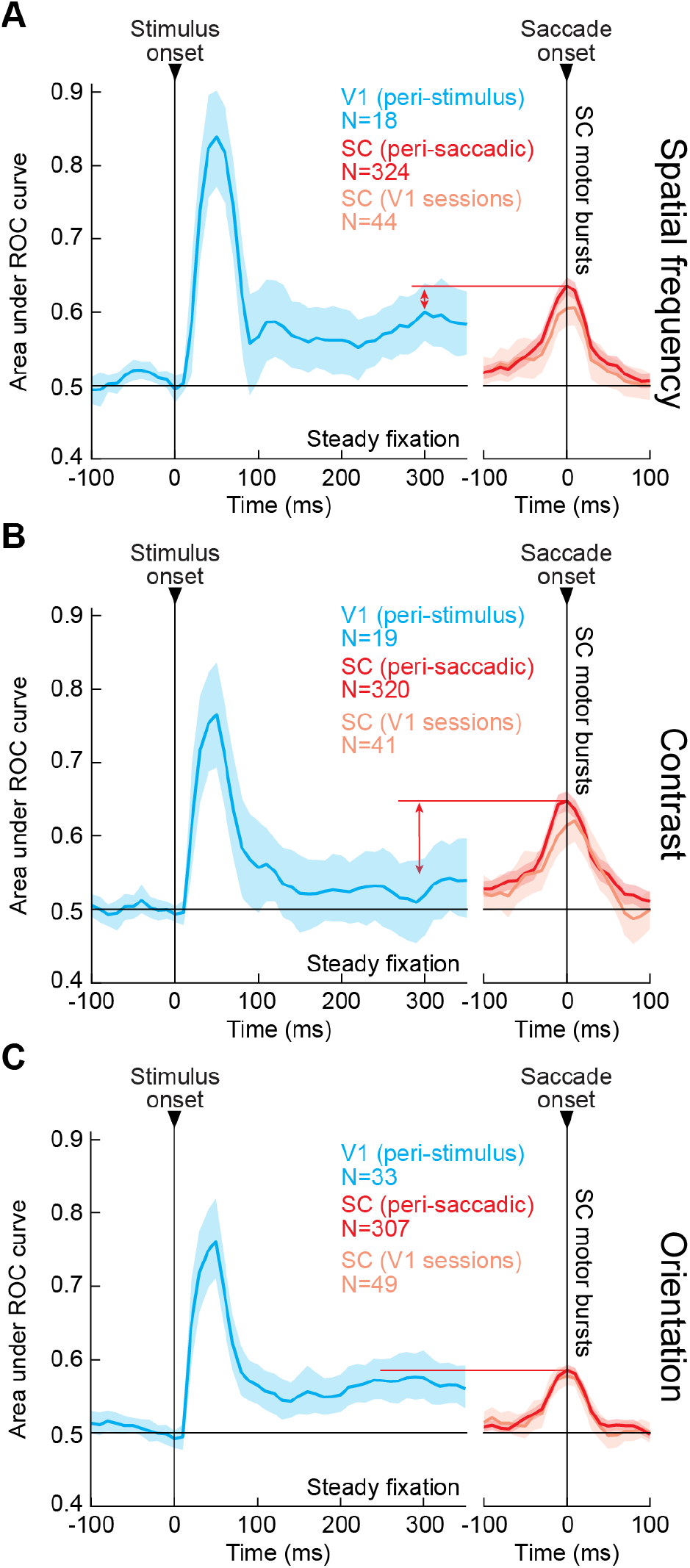
SC motor bursts as a potential means for providing a trans-saccadic sensory bridge of visual information across rapid eye movements. **(A)** We simultaneously recorded primary visual cortex (V1) activity with some of our SC neurons (with overlapping RF locations; Methods). For each V1 neuron, we calculated peri-stimulus AUC discrimination performance between its most and least preferred images in the visual burst epoch (light blue). After an initial peak in the stimulus-evoked visual bursts, the visual representation in peripheral V1 returned to a lower, but still feature-tuned, baseline during steady-state gaze fixation, like in V4^20,45^. The sensory-tuned peri-saccadic SC motor burst signal that we observed (red: all neurons; orange: simultaneously recorded neurons with V1) had AUC discrimination performance (between most and least preferred images) that peaked to a level similar to how V1 discriminated peripheral images in steady-state fixation. Thus, when the eyes rapidly move and cortex expects uncertain retinal input, the SC still possesses a sensory representation of the peripheral saccade-target appearance (also see Fig. 3G-I). **(B, C)** Similar results for the contrast and orientation image manipulations. In all cases, peak peri-saccadic AUC discrimination performance in the SC was higher than or similar to steady-state AUC discrimination performance of peripheral image information in the visual cortex during fixation. Error bars: 95% confidence intervals.

Thus, during saccades, the SC possesses a reliable sensory representation, which may provide a trans-saccadic sensory bridge for perception, exactly when visual sensory brain areas might lack coherent retinal image input due to rapid eyeball rotations.

### Link to peri-saccadic perception

The results of Fig. 6, coupled with dissociation of the SC motor burst modulations from movement properties (e.g. Fig. 2 and refs. ^12,13^), suggest that sensory tuning in SC neuronal movement commands may be more relevant for the SC’s ascending pathways to the cortex^32,33^ than for its descending projections to the oculomotor control network. While such ascending pathways have historically been suggested to provide only the vector information of the saccade, to spatially remap retinotopic visual representations^31,32,34^, sensory tuning in SC motor bursts could allow the same pathways to additionally relay a sensory prediction signal of the peripheral saccade target appearance; such a peripheral preview (even if it is coarser than foveal visual analysis) could be perceptually useful, for example, in predicting the foveal visual sensory consequences of saccades^35^, or in bridging trans-saccadic perception^36-38^. It could also pre-saccadically enhance perception of the saccade target^22-25^ and aid in visual search^39^.

If so, then classic peri-saccadic perceptual phenomena, such as saccadic suppression^40^ and mislocalization^41^, which are thought to depend on SC-sourced corollary discharge signals^33,34,42,43^, should also be expected to vary with different saccade targets. In other words, if similar saccades to different images are associated with different SC motor bursts, and if these bursts relay more than just the vector of the saccades via corollary discharge, then peri-saccadic phenomena dependent on SC-sourced corollary discharge should also depend on saccade target appearance.

We recently investigated perceptual saccadic suppression with saccades made across backgrounds of low or high spatial frequency^44^. Saccadic suppression was significantly stronger for the low spatial frequency condition^44^. The saccade targets for the different background images were indeed different images, exactly like in our neurophysiological manipulations here. Therefore, it is indeed plausible that sensory tuning in the SC motor bursts might be linked to modulating the properties of peri-saccadic suppression.

To explicitly test this, we replicated the current study’s paradigms in human perceptual experiments probing both peri-saccadic suppression and peri-saccadic mislocalization. We asked humans to generate saccades towards a localized low or high spatial frequency grating as the saccade target (Supp. Fig. 10A; Methods). Given that low and high spatial frequencies were differentially represented in our neurons (Fig. 3D, saccade epoch), and given our recent results with low and high spatial frequency backgrounds^44^, we predicted that peri-saccadic perceptual performance will vary across these two conditions.

In the suppression paradigm, we peri-saccadically flashed brief, low contrast probes, and we measured perceptual contrast sensitivity curves^44^. Supp. Fig. 10B shows peri-saccadic probe times, and Supp. Fig. 10C shows an example subject’s eye movement properties for the two different target images. We ensured matched saccade vectors and kinematics across all experiments (Methods). In Supp. Fig. 10D, the subject’s perceptual performance is shown for either intra-saccadic flashes (red and blue) or post-movement ones at perceptual recovery (yellow and light blue). At perceptual recovery (post-movement probes), probe visibility was not affected by the saccade target appearance; the psychometric curves were the same whether the saccade target was low or high in spatial frequency. This confirms that visual processing of the probes was not modulated a priori, perhaps via long-range lateral interactions, by the mere visual appearance of the saccade target. Peri-saccadically (when SC motor bursts differentially represented saccade target appearance; Figs. 1-3), saccadic suppression was stronger (i.e. there were higher perceptual thresholds) for low spatial frequency targets. These results held across the population (Supp. Fig. 10E). Thus, the visual appearance of the saccade target did indeed matter for the strength of peri-saccadic sensitivity.

Our mislocalization paradigm was the same, except that the probes were always high in contrast, and the subjects had to point to where they saw them (Methods). Again, we found stronger peri-saccadic mislocalization for a low spatial frequency saccade target compared to a high spatial frequency one (Supp. Fig. 10G-I), and this difference decreased for probes farther away in time from saccade execution (Supp. Fig. 10J).

Therefore, peri-saccadic perceptual phenomena that are believed to depend on ascending SC projections to the cortex^33,34,42,43^ also depend on the visual appearance of the saccade target, suggesting that sensory tuning in SC neuronal movement commands may provide more than just the vector information of executed movements via SC-sourced corollary discharge signals.

## Discussion

We observed a robust visual sensory representation embedded within SC neuronal eye movement commands (e.g. Figs. 1, 2). This representation amplifies weak visual signals (e.g. Fig. 3D-F), and it is strong for images of real-life objects (e.g. Fig. 4). It also occurs at a time in which a retinal transfer of visual information about the saccade target ballistically takes place from the periphery to the fovea, causing high afferent sensory uncertainty.

Given that the SC integrates a large amount of visual information from the retina and cortex^46,47^, sensory tuning in its saccade movement commands is an ideal means for internally maintaining evidence about the visual appearance of saccade targets, at a time when such evidence from external afference might be unreliable. Indeed, amplification of weak visual signals during motor bursts fits with a large range of evidence that perception is enhanced at the saccade target around the time of eye movements^22-25^. A pre-saccadic strengthening of feature-tuned neuronal representations in the visual cortex also takes place^45^. Finally, our perceptual peri-saccadic suppression and compression results are consistent with a potential role of SC neuronal movement commands in relaying more than just the vector of intended eye movements to the rest of the visual system. Interestingly, even though SC visual responses to small spots of light are weak in the retinotopic lower visual field, SC motor bursts for downward saccades to such spots are stronger^12^. Thus, amplification of weak SC visual signals might be a general property of this structure’s motor bursts.

The SC still contributes to saccade generation. However, even though it is often suggested that SC motor bursts are critical for saccade control^8,10,11^, increasing evidence suggests a significantly smaller role, consistent with our observations. For example, SC motor bursts are affected by audio-visual combinations without altered saccades^48^. Moreover, saccades to a blank are often associated with opposite changes in SC bursts and movement kinematics, relative to what happens with saccades to a visible spot^12,13^. Finally, reward expectation alters all aspects of SC responses in saccade paradigms, including both visual and motor bursts^16^. Therefore, SC movement commands can be useful for other aspects of active behavior. Indeed, given that the SC is causally necessary for maintaining visual object representations in a patch of the temporal cortex^49^, and given the sensitivity of SC neuronal movement commands to images of coherent visual objects (Fig. 4), it is very intriguing to consider the possibility that the SC can influence the visual temporal cortex also at the time of saccades. This would provide an ideal active vision pathway to the processing of natural visual scenes in normal behavior.

Looking forward, our results can help resolve further long-standing mysteries about SC neuronal movement commands. For example, population activity spreads across the SC during saccades^50^, but no convincing explanation for this exists. Since small spot targets are spectrally broadband stimuli, it could be that spreading activity is just a manifestation of different spatio-temporal mRF properties for images of different spatial frequencies. This, along with investigating whether the SC can causally influence cortical visual feature updating across saccades, will undoubtedly significantly clarify long-lasting mysteries on perception, action, and the self-monitoring internal processes that necessarily link the two.

## Acknowledgements

We were funded by the Deutsche Forschungsgemeinschaft (DFG): (1) BO5681/1-1; and (2) SFB 1233, Robust Vision: Inference Principles and Neural Mechanisms, TP 11, project number: 276693517. We were also funded by the Medical Faculty of the University of Tübingen.

## Author contributions

MPB, ARB, AFD, ZMH performed neurophysiological experiments. MPB, AFD, ZMH performed psychophysical experiments. MPB, ARB, AFD, ZMH analyzed data and wrote manuscript.

## Data availability

The databases generated during the current study are available from the corresponding author on reasonable request.

## Supplementary figures

**Supp. Fig. 1.**
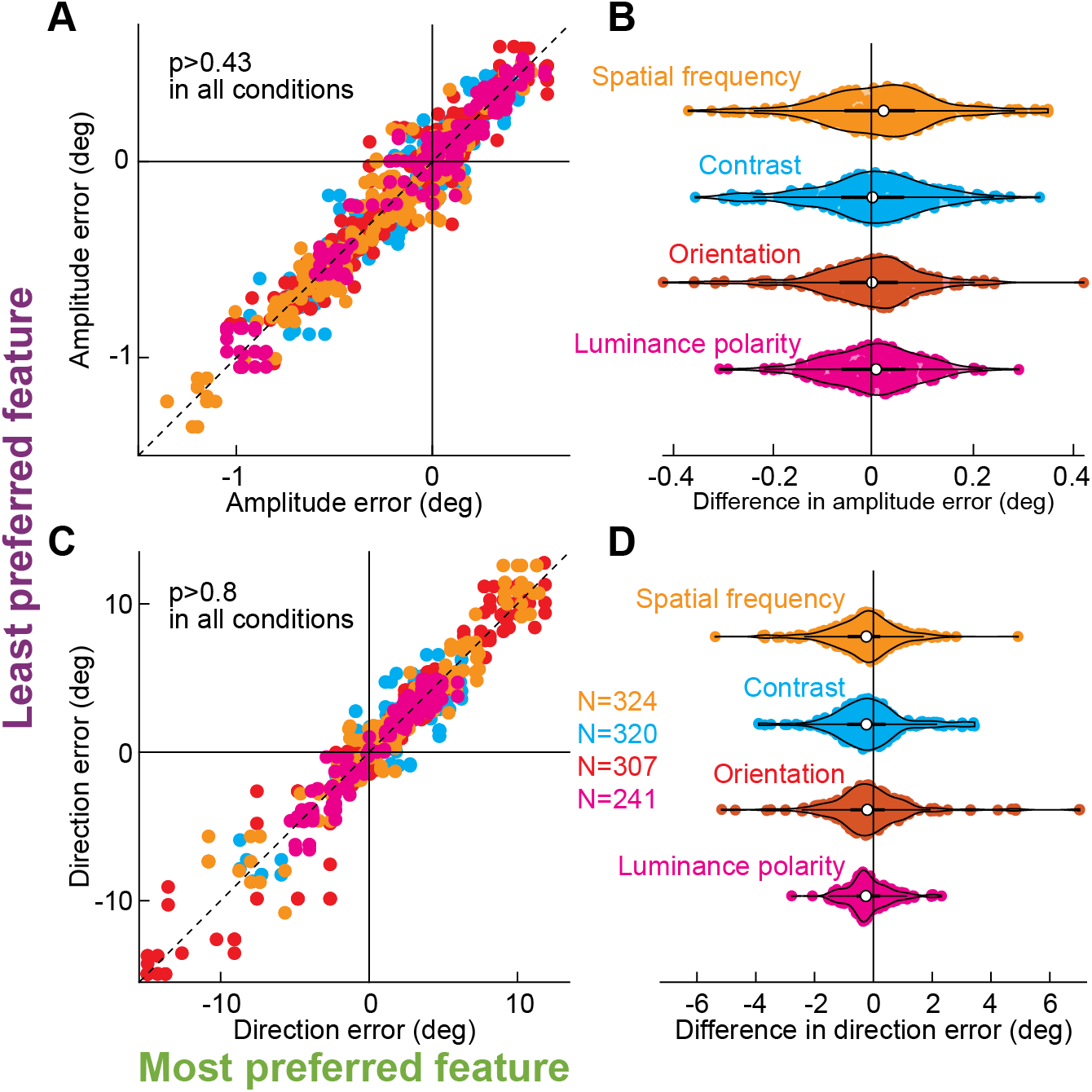
Independence of sensory tuning in SC neuronal movement commands from eye movement metric properties. **(A)** For each image manipulation of Fig. 2 (different colors), and for each neuron (individual symbols), we plotted the amplitude error of the saccades to the most and least preferred images of the neuron (based on its motor burst strengths). Despite the large differences in saccade motor bursts (Fig. 2), the amplitude errors were similar across images. This confirms our vector matching procedures (Methods). P-values indicate rank-sum test results from each image manipulation. **(B)** Distributions of amplitude error differences between most and least preferred image trials from **A**. The violin plots always straddled zero. **(C, D)** Similar observations with saccade direction errors. The numbers of neurons indicated apply to **A, B** as well.

**Supp. Fig. 2.**
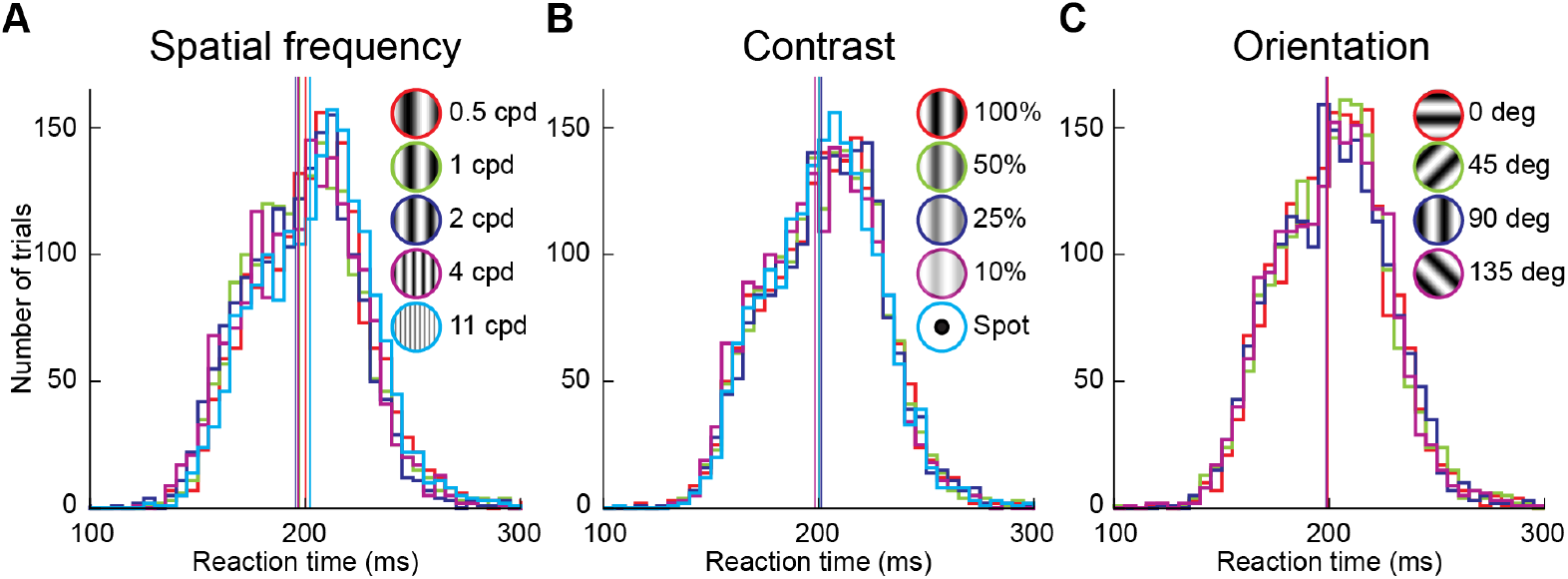
Independence of sensory tuning in SC neuronal movement commands from intrinsic image salience, as inferred from saccadic reaction times. **(A-C)** In each of the spatial frequency, contrast, and orientation image manipulations, we used a delayed-saccade paradigm to enforce a period of steady-state gaze fixation between image onset and saccade triggering. This allowed us, as much as possible, to equalize saccadic reaction times across image features within each image manipulation, as can be confirmed from the strongly overlapping saccadic reaction time histograms in each panel. Thus, even though some image features, like low spatial frequencies^17^, might be more intrinsically salient than others we equalized this as much as possible by our delayed-saccade paradigm. Note also that in our luminance polarity image manipulation, we used reflexive saccades instead, and the results were unchanged despite strong differences in saccadic reaction times with different image features (see Supp. Fig. 3). Vertical lines indicate mean reaction times for each image feature.

**Supp. Fig. 3.**
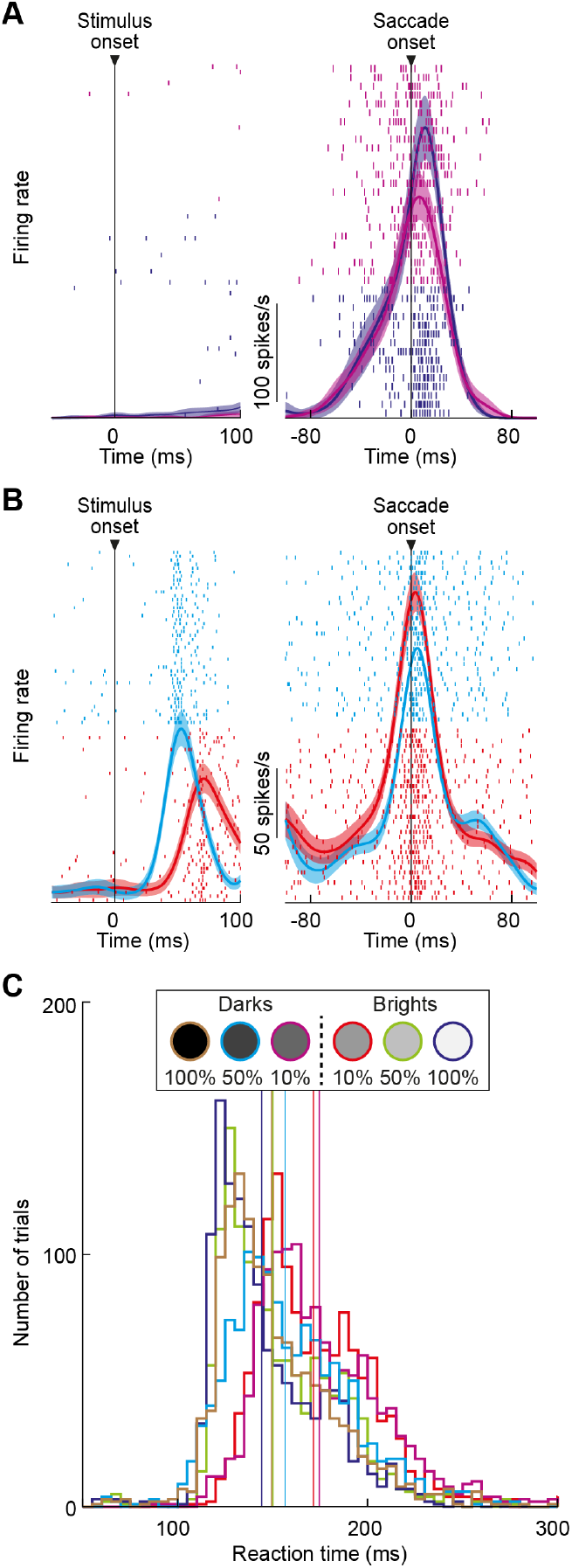
Independence of sensory tuning in SC neuronal movement commands from reflexive versus delayed saccades. **(A)** An example neuron’s firing rates from our luminance polarity image manipulation. In this image manipulation, we avoided delayed saccades, and the monkeys reflexively looked at the peripheral stimulus as soon as it appeared. This example neuron had almost non-existent visual responses to stimulus onset, but it had strong motor bursts. For two different image features, the motor bursts were very different, similar to the example neuron of Fig. 1. Error bars: 95% confidence intervals, and the numbers of trials are indicated by the number of spike raster rows shown. Other conventions are similar to Fig. 1, and the colors indicate the individual image features, as per the legend in **C. (B)** A second example neuron possessing both visual and motor bursts. Note how the visual burst had strongly different latencies from stimulus onset in the two shown conditions, which was also reflected in different saccadic reaction times (**C**). Nonetheless, the motor bursts were still sensory-tuned like in the delayed-saccade paradigm of Fig. 1. Also note how the neuron flipped its image feature preference between visual and motor burst epochs, showing weaker visual bursts but stronger motor bursts for the same image. This is consistent with a transformed SC representation of images at the time of saccade triggering (also see Fig. 3 and Supp. Fig. 6). Error bars: 95% confidence intervals, and trial numbers can be inferred from the shown spike rasters. **(C)** With the reflexive saccade paradigm used in this image manipulation, saccadic reaction times strongly depended on image contrast, and there were modulatory effects of luminance polarity^19^. Thus, whether saccadic reaction times were equalized (Supp. Fig. 2) or not (this figure), sensory tuning in SC neuronal movement commands was still robustly observed. Vertical lines indicate mean saccadic reaction times for each image feature.

**Supp. Fig. 4.**
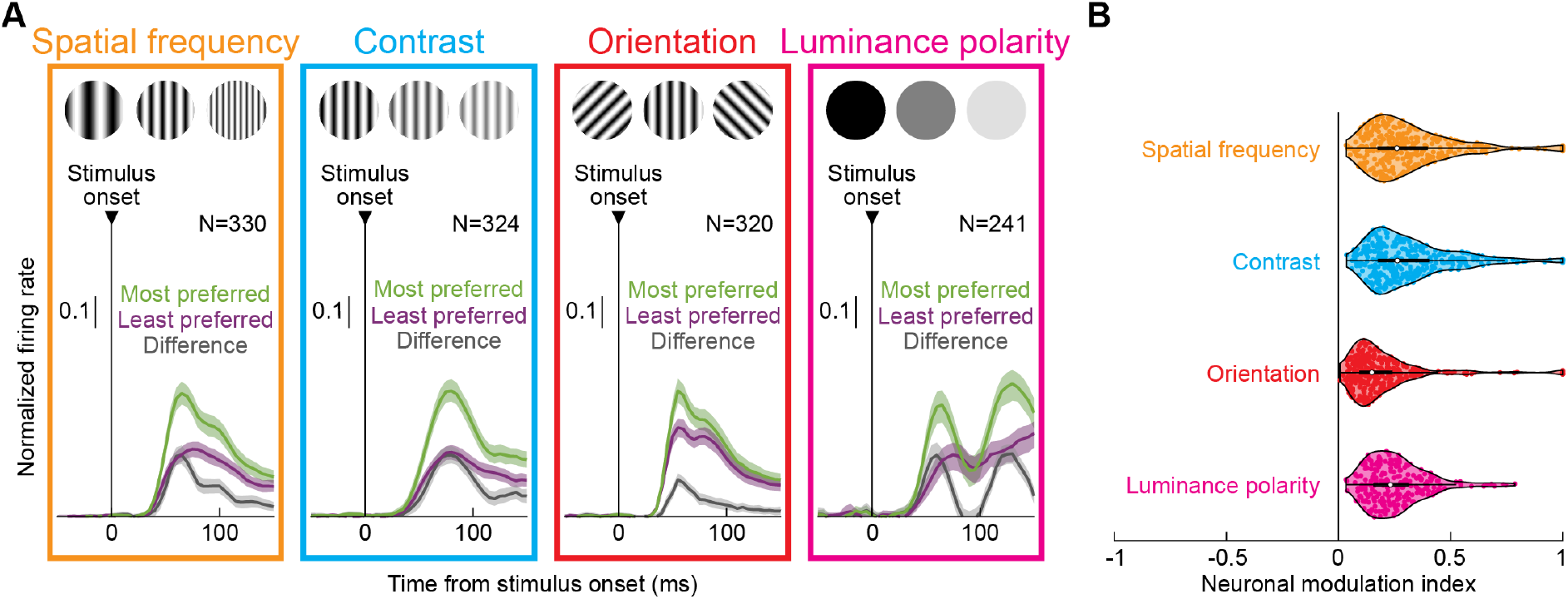
Sensory tuning in SC visual responses. **(A)** Analyses like those in Fig. 2A, but for the visual responses of the neurons rather than their activity at the time of saccade triggering. The differences in firing rates between most and least preferred images in the visual bursts were smaller than in the saccade bursts of Fig. 2A, consistent with the AUC discrimination performance results documented in Fig. 3A-C. **(B)** Neuronal modulation indices from the visual burst epoch; these were calculated similarly to the modulation indices in the motor bursts, but based on visual burst measurements and feature preferences (Methods). All conventions are similar to Fig. 2. Also note that the luminance polarity image manipulation was from the reflexive saccade paradigm. Therefore, there were secondary elevations in firing rates after the initial visual responses in **A**, reflecting the saccade motor bursts. Error bars: 95% confidence intervals, and neuron numbers are indicated in **A**.

**Supp. Fig. 5.**
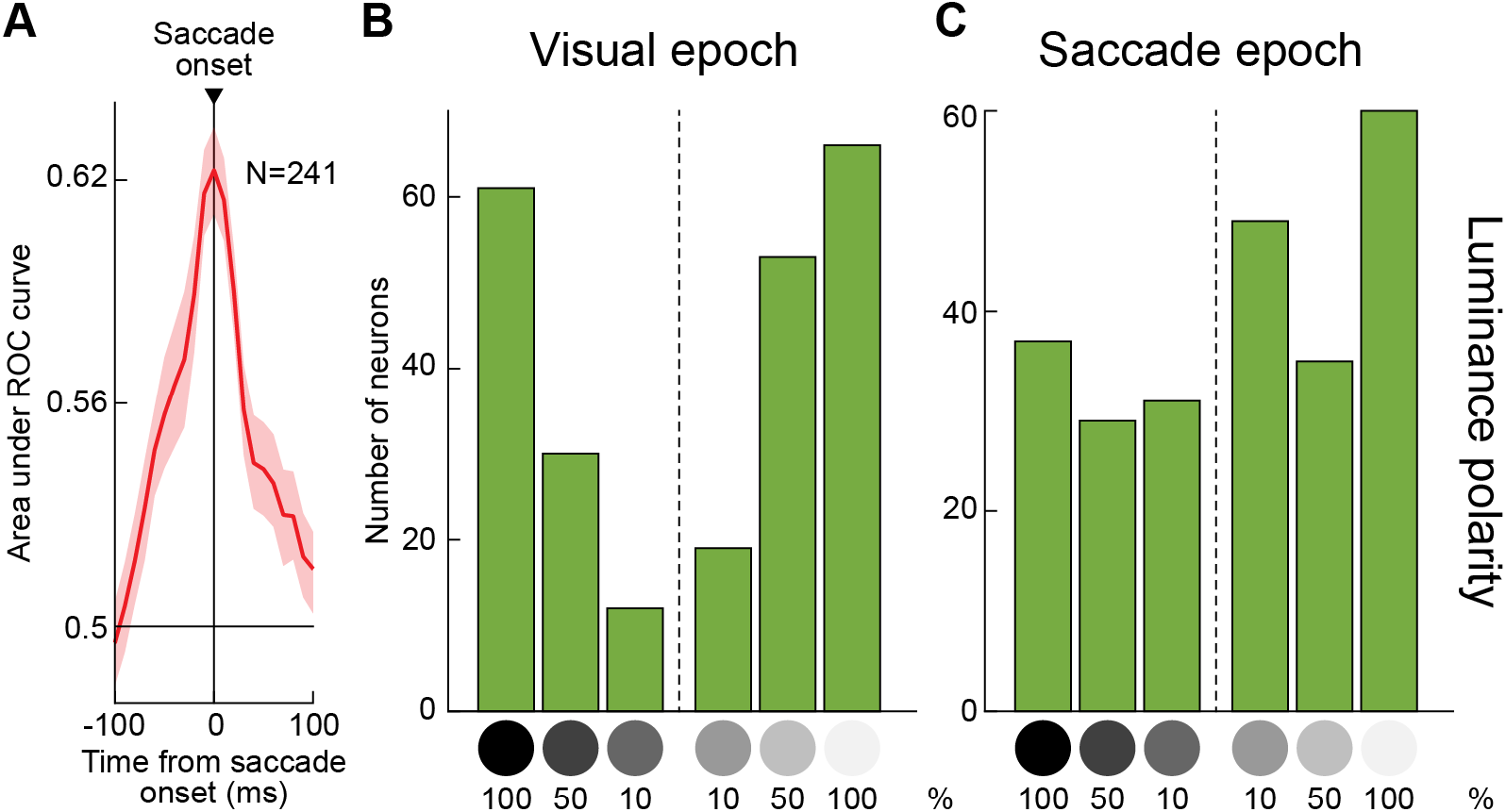
Similar observations to Fig. 3A-F during the reflexive saccade paradigm. **(A)** From the luminance polarity image manipulation, we plotted peri-saccadic AUC discrimination performance across neurons. Consistent with Fig. 3 and the example neurons of Supp. Fig. 3, there was a peak in AUC discrimination performance at the time of SC motor bursts. **(B, C)** Also consistent with Fig. 3, the distribution of preferred image features at saccade onset (**C**) was broader than that at stimulus onset (**B**), suggesting amplification of weak visual signals at the time of saccade generation even with reflexive, visually-guided saccades. Supp. Fig. 3B shows an example neuron with such amplification at the time of saccades. Also see Supp. Fig. 7 for high-dimensional population activity trajectories in the reflexive saccade paradigm.

**Supp. Fig. 6.**
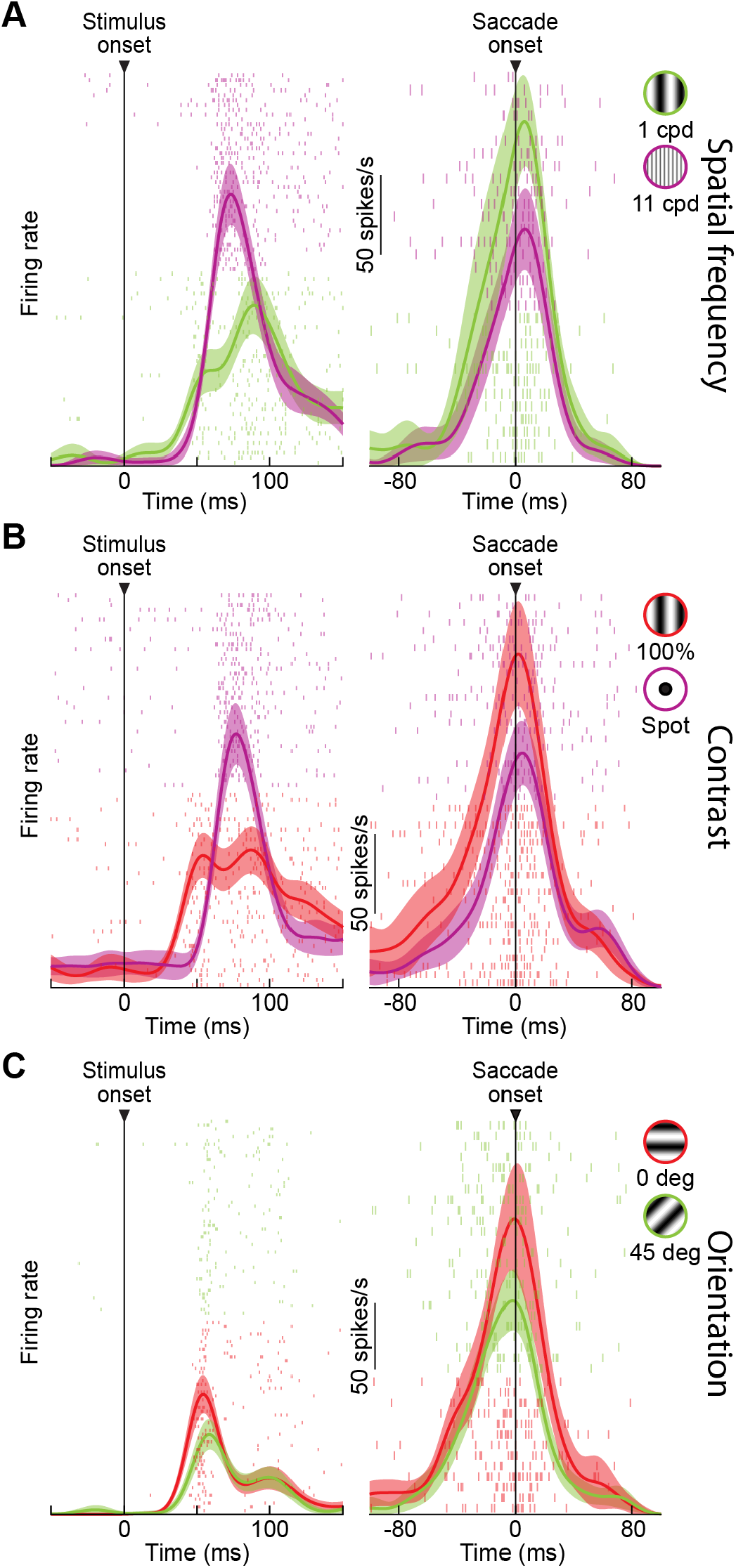
Potential transformation of image preferences between stimulus and saccade onsets in individual SC neurons. **(A-C)** Three example neurons from our three image feature manipulations with the delayed saccade paradigm, demonstrating how a changed feature preference can occur between visual and motor epochs with simple grating stimuli. Note how the weak signals in the visual epochs in **A, B** were transformed into stronger motor bursts at the time of saccade triggering. Also note that this is similar to the example neuron of Supp. Fig. 3B in the immediate, reflexive saccade paradigm. The neuron in **C**, on the other hand, did not exhibit an altered preference between its visual and motor epochs. Error bars: 95% confidence intervals, and trial numbers can be inferred from the shown spike rasters

**Supp. Fig. 7.**
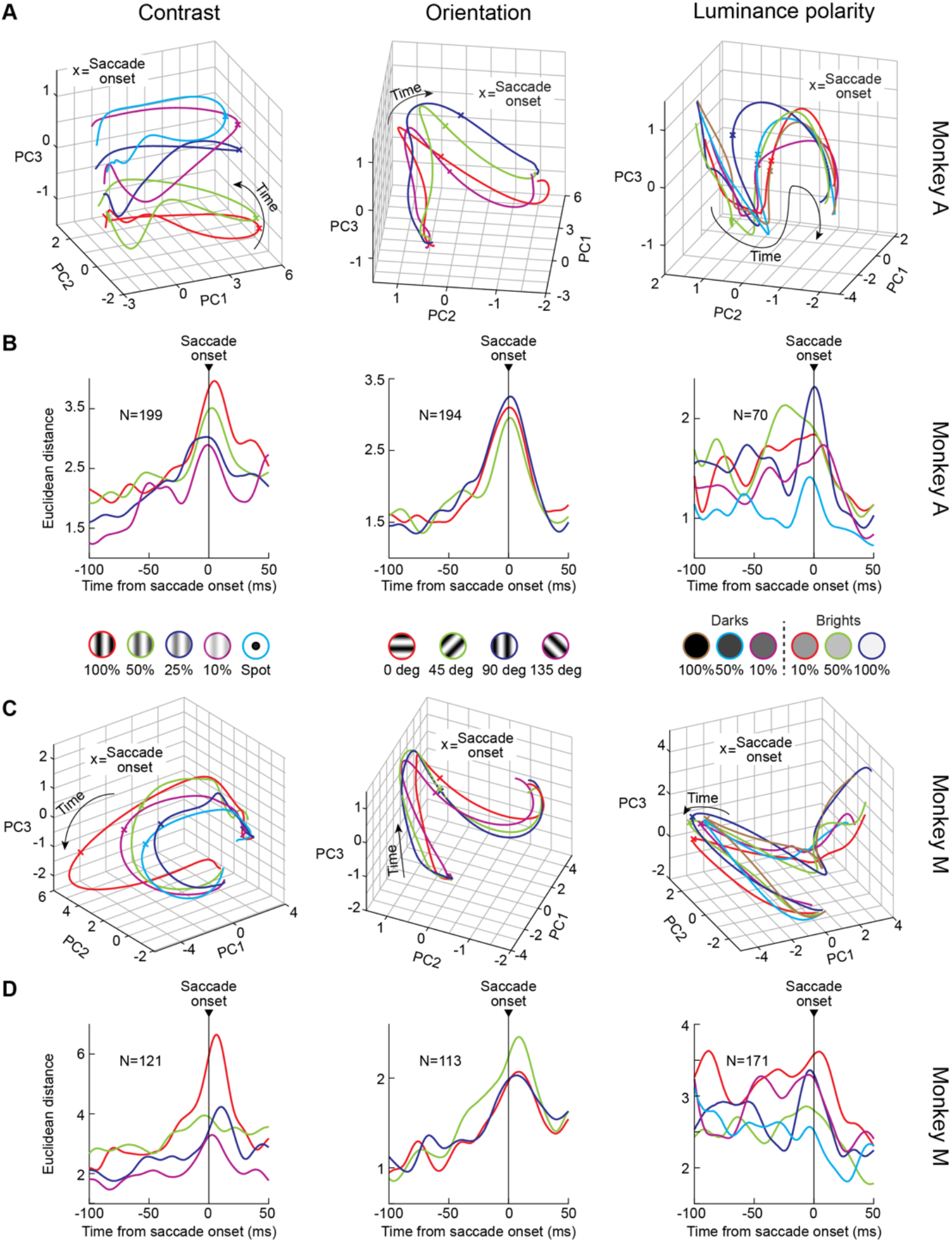
Embedding of image feature information in SC population activity at the time of saccades. **(A)** Monkey A population activity trajectories in the first 3 principal components after PCA decomposition in the contrast, orientation, and luminance polarity image manipulations (spatial frequency was shown in Fig. 3H). Consistent with Fig. 3H, SC neurons occupied different manifolds in population activity space at the time of saccade triggering for different image features. Note that in luminance polarity, the saccades were reflexive. Thus, visual and motor bursts occurred in close temporal proximity to each other. Nonetheless, their transformation into quasi-orthogonal manifolds between the visual and motor epochs was still visible, consistent with Fig. 3G. **(B)** For each image manipulation, we picked a reference condition (spot for contrast, 135 deg for orientation, and 100% dark for luminance polarity), and we then plotted the peri-saccadic Euclidean distance of high-dimensional SC population activity from this condition at the time of saccade generation. In each case, the Euclidean distances were different for different image features, suggesting the embedding of sensory information at the time of SC motor bursts (despite vector and kinematic saccade matching). Note that for luminance polarity, the sustained elevation of Euclidean distances before saccade onset reflects the visual epochs of this reflexive saccade task, which were also sensory-tuned. **(C, D)** Similar results from monkey M. Also see Supp. Fig. 8D for consistent results of sensory tuning in SC neuronal movement commands with real-life object images.

**Supp. Fig. 8.**
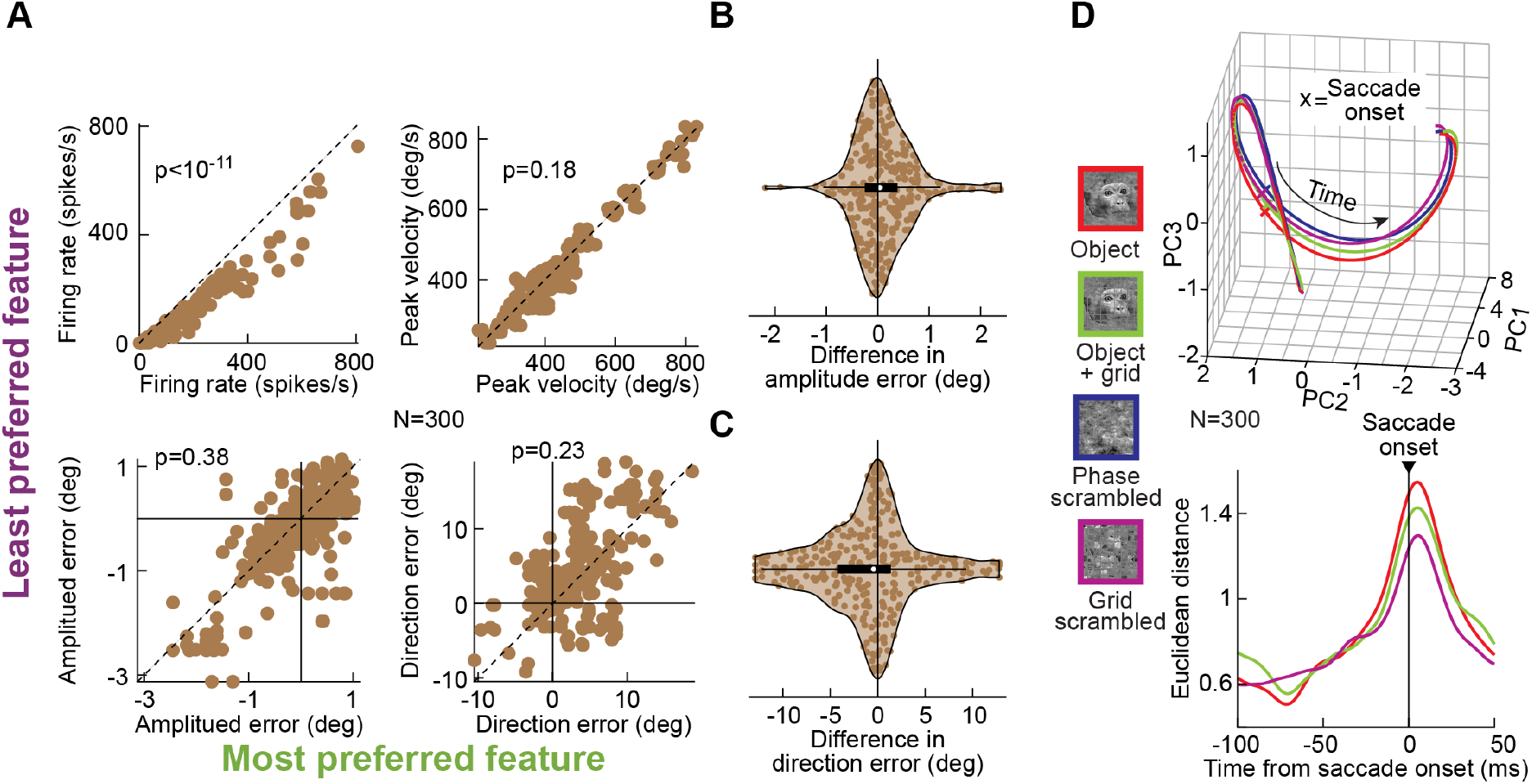
Real-life object representations in SC neuronal movement commands. **(A)** Plots similar to Fig. 2B, C and Supp. Fig. 1A, C showing a dissociation between motor burst effects between most and least preferred images (top left) and saccade kinematics (top right) or saccade metrics (bottom left and right) in the experiment testing real-life object images (Fig. 4). **(B)** Distributions of saccade amplitude and direction error differences between most and least preferred images like in Supp. Fig. 1B, D, consistent with the interpretation that SC motor burst differences in this experiment were not explained by systematic differences in eye movement parameters (also see Fig. 4D, E). **(D)** Peri-saccadic PCA-space population trajectories (top) and high-dimensional space Euclidean distances (bottom) from both monkeys in the experiments with object images. Note how object and object+grid images (having coherent visual form images within them) were more differentiated from phase-scrambled images (higher Euclidean distances) at the time of saccade triggering than grid-scrambled images, consistent with Fig. 4F.

**Supp. Fig. 9.**
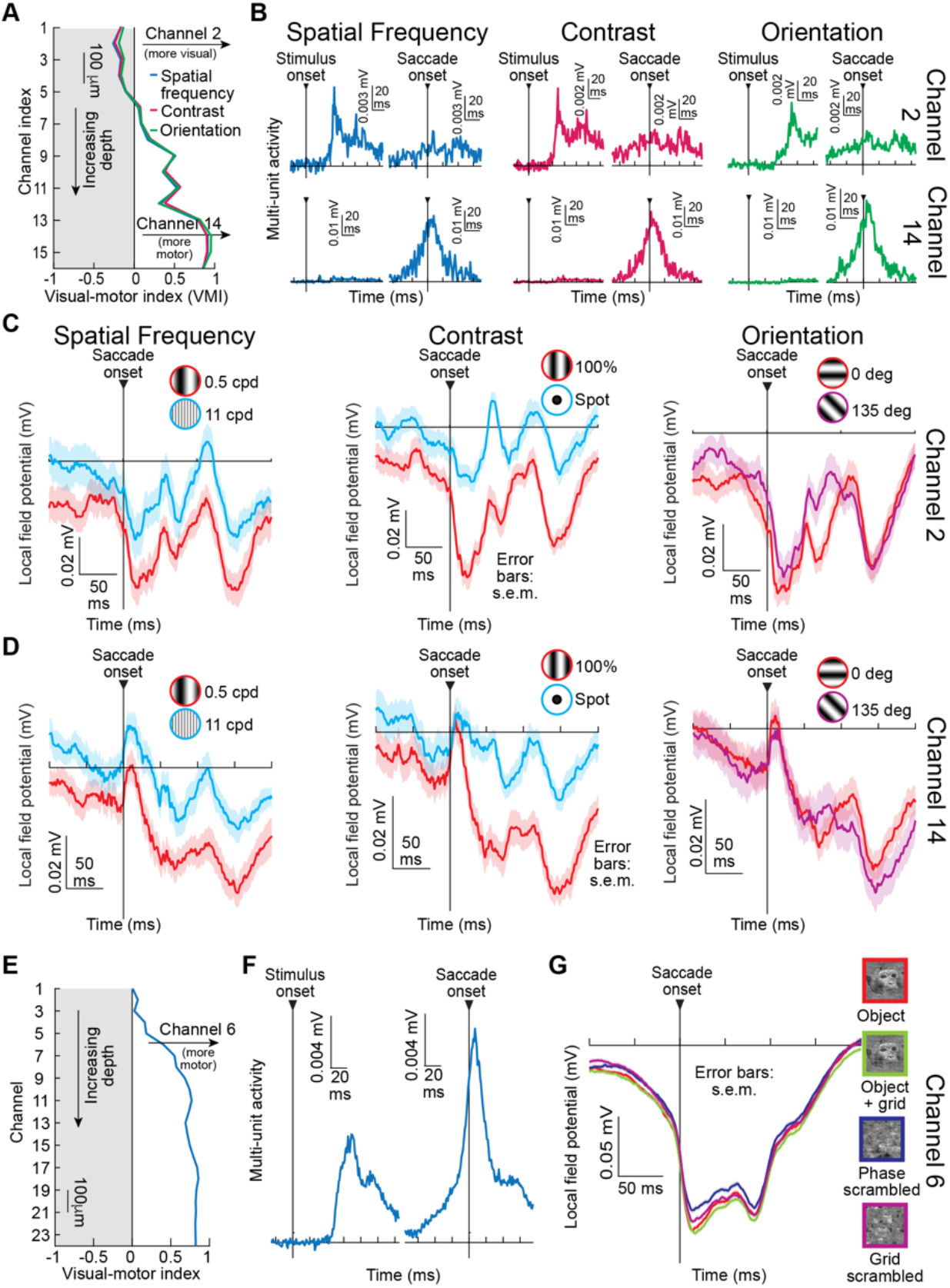
Embedding of sensory information at saccade onset within the deeper SC layers. **(A)** We calculated a visual-motor index (VMI) (Methods and ref. ^51^) across electrode depths from the example session shown in Fig. 1. The VMI, which is inferred from multi-unit activity (MUA) near a given electrode contact, is positive for more motor layers and negative for more visual layers, and the example neuron of Fig. 1 was recorded from channel 14 (that is, from a strongly motor layer). The VMI was calculated for each image manipulation separately (3 colors), and it was robust across them. **(B)** Example MUA activity profiles near stimulus or saccade onset from the same example session. Responses are shown from channel 2 and channel 14, demonstrating how channel 2 was predominantly visual (no motor bursts) and channel 14 was predominantly motor (no visual bursts). **(C)** Local field potential (LFP) profiles around saccade onset for two example features (e.g. 0.5 and 11 cpd) from each image manipulation tested in this session (spatial frequency, contrast, and orientation). The LFP responses from channel 2 are shown. There were differences in peri-saccadic LFP responses for different image features of the saccade targets, despite matched saccade kinematics and metrics. This is consistent with the presence of sensory information in the local SC network at the time of saccade motor burst generation. Note how the effect was weakest for the orientation image manipulation. **(D)** Similar analyses from the deeper motor layer of channel 14 (where the example neuron of Fig. 1 was recorded). There was still a clear sensory signal in the peri-saccadic LFP responses despite the depth of the recording, consistent with the results of Figs. 1-3, 5. Again, the effect was weakest in the orientation image manipulation. **(E-G)** VMI, MUA, and LFP responses from another session from the real-life object experiment. Peri-saccadic LFP responses from an example motor layer (same as that of the example neuron of Fig. 4B) also differentiated between coherent and scrambled object images (**G**). Error bars in all cases: s.e.m.

**Supp. Fig. 10.**
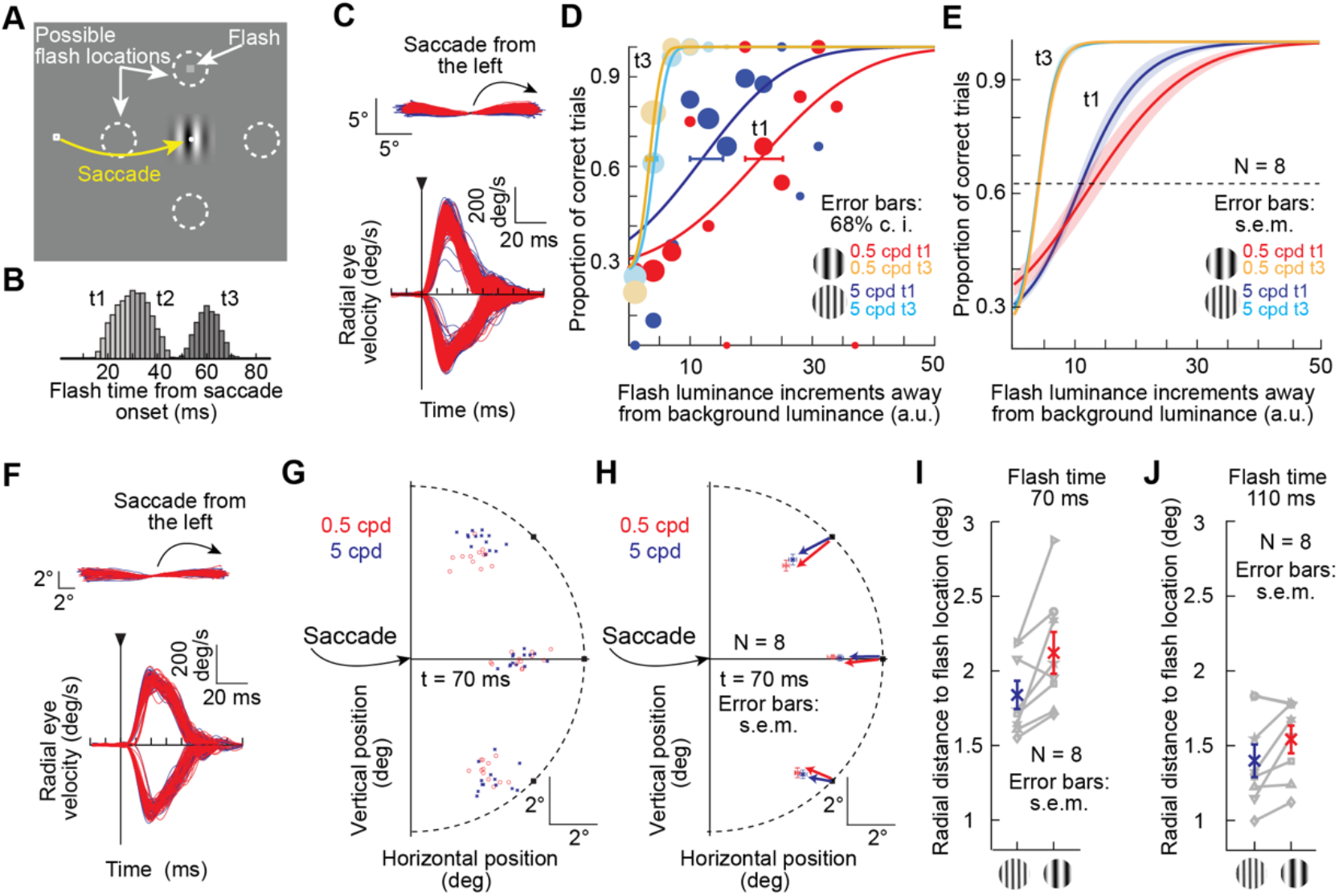
Dependence of peri-saccadic perceptual suppression and mislocalization on the visual appearance of the saccade target. **(A)** To test if a sensory signal in SC neuronal movement commands could be relevant for phenomena believed to depend on SC-sourced corollary discharge, we asked human subjects to generate saccades to low or high spatial frequency gratings. At a random time relative to saccade onset, a probe flash was presented briefly at one of four locations around the saccade target. **(B)** Distribution of probe flash times relative to saccade onset across all of our experiments. We grouped flash times into three groups, with t1 being expected to be associated with maximal perceptual suppression and t3 being associated with perceptual recovery. **(C)** Example saccade trajectory and radial eye velocity profiles from one subject, demonstrating matched metrics and kinematics for the two different tested spatial frequencies as the saccade targets (Methods). **(D)** Perceptual performance as a function of probe flash luminance from the same subject at t1 and t3. Despite the matched saccades, the subjects’ perceptual threshold at t1 (which was strongly elevated relative to t3 as expected) was higher for the low spatial frequency saccade target than for the high spatial frequency saccade target. Thus, at maximal suppression, the visual appearance of the saccade target influenced the strength of peri-saccadic perceptual suppression (a phenomenon that is believed to depend, at least in part, on SC-sourced corollary discharge^33^). Error bars: 68% confidence intervals. **(E)** Similar observations across all subjects. Note how perceptual thresholds at t3 were similar for the two different saccade target appearances, suggesting that the differences at t1 were not merely due to visual-visual interactions between gratings and probe flashes. Also note that t2 performance, which is not shown for clarity, fell in between that in t1 and t3, as expected from the known time course of perceptual saccadic suppression. Error bars: s.e.m. **(F)** Eye movement parameters from the same subject as in **C**, but when the subject performed our second variant of the experiment, probing peri-saccadic perceptual mislocalization (Methods). **(G)** Despite matched saccades, the subject mislocalized high contrast peri-saccadic flashes more when the saccade target was a low spatial frequency (compare red click locations to blue click locations; black squares show the actual flash locations); example data subset shown. Thus, the well-known mislocalization phenomenon depends on the visual appearance of the saccade target. **(H)** Summary of median click locations in the two conditions across subjects (mean and s.e.m. across 8 subjects shown). **(I)** For peri-saccadic flashes, we plotted the mislocalization error as the radial distance from subjects’ click locations to actual probe flash locations. Individual subject distances (based on median click locations) are shown in gray, and the blue and red symbols show the mean and s.e.m. across subjects. There was larger mislocalization for the low spatial frequency saccade targets (all but one subject showed the effect). **(J)** At longer times after saccade onset, there was perceptual recovery and a smaller difference in mislocalization between low and high spatial frequency saccade targets.

## Methods

### Experimental animals and ethical approvals

All animal experiments were approved by ethics committees at the regional governmental offices of the city of Tübingen, Germany (*Regierungspräsidium Tübingen*).

We recorded superior colliculus (SC) neuronal activity from two adult, male rhesus macaque monkeys (M and A), aged 8-9 years and weighing 9.5-10 kg. We also recorded primary visual cortex (V1) neuronal activity, simultaneously with SC activity, from monkey A.

### Animal laboratory setup and animal preparation

The animal laboratory setup was the same as that described in our recent work^30^. Briefly, each animal was placed in a darkened room ∼72 cm from a calibrated and linearized cathode-ray-tube (CRT) display (spanning approximately 30 deg horizontally and +/-23 deg vertically). We controlled the stimulus presentations and data acquisition procedures using a custom-built modification of PLDAPS^52^, interfacing with the Psychophysics Toolbox^53-55^ and an OmniPlex data acquisition system (Plexon, inc.).

We prepared the animals for behavioral training and neurophysiological experiments earlier^30,56,57^. Briefly, in each animal, we implanted a head holder, to stabilize head position, and a scleral search coil in one eye^58^, to allow tracking eye movements with high quality using the electromagnetic induction technique^59^. We also implanted a recording chamber centered on the midline and tilted 38 deg posterior of vertical. In monkey A, we positioned the chamber to allow access to the SC in the upper half of the chamber and dorsal portions of V1 in the lower half. SC and V1 recordings were performed in the left hemisphere of monkey A. Monkey M’s SC recordings were performed in both hemispheres.

We recorded neuronal activity using 16- or 24-channel linear electrode arrays with 50 μm inter-electrode spacing (V-Probes from Plexon, inc.).

### Animal behavioral tasks

Our primary behavioral task was the “Saccades-to-X” paradigm. This was a modified version of the classic delayed, visually-guided saccade task, with the main difference being that we used an image as the eccentric saccade target rather than just a small spot of light.

Each trial started with the presentation of a white fixation spot (10.8 by 10.8 min arc dimensions) presented at display center. The spot had a luminance of 79.9 cd/m^2^, and it was presented over a gray background of luminance 26.11 cd/m^2^. After the monkey fixated the spot stably for 300-700 ms, an image was presented at an eccentric location. The monkey was instructed to maintain fixation on the spot even after the onset of the eccentric image. After 500-1000 ms of successful gaze fixation, the central fixation spot was removed, instructing the monkey to generate a saccade towards the center of the eccentric image. To minimize saccade vector variability across trials, which was critical for ruling out metric and kinematic changes in the saccades as the main sources of our results, we always provided a clear visual marker at the center of the eccentric image, which served as an anchor for directing the saccades towards. This marker was superimposed on the eccentric image, and it consisted of a white spot, just like the fixation spot, surrounded by a gray disc (0.54 deg diameter) of the same luminance as the background. The surrounding gray disc ensured that the marker spot was visible irrespective of the background image, allowing us to experimentally control, as much as possible, trial-to-trial variability of the saccades made to the extended images. This was important because saccades can span a range of locations on an extended foveated image^60^, complicating the interpretation of whether motor burst changes were due to the image or due to different saccade vectors activating different parts of movement-related response fields (mRF’s). In subsequent post-hoc analyses (see below), we further controlled trial-to-trial variability of the saccades, again to rule out a trivial motor variance explanation of our results. The monkey was rewarded for successfully generating a saccade towards the image center within 500 ms from fixation spot offset, as well as for maintaining gaze on the eccentric image center for an additional 500 ms.

In different blocks of trials, we used different series of images as the eccentric saccade targets. For example, in Saccades-to-Spatial-Frequency, the saccade target image consisted of a disc of 3 deg radius, the inside of which was a vertical, stationary sine wave grating of 100% contrast and a specific spatial frequency. The spatial frequency was randomly picked for each trial from among the following values: 0.5, 1, 2, 4, and 11 cycles/deg (cpd).

In the Saccades-to-Contrast variant of the task, the grating image was still vertical like in Saccades-to-Spatial-Frequency, but it now had a fixed spatial frequency (1 cpd). Across trials, we varied the contrast of the grating from among the following values: 100%, 50%, 25%, 10%, and 0%. Note that the 0% contrast grating had no luminance variation in it at all, but the marker spot (described above) was still visible at its center. Therefore, the 0% contrast condition looked identical to a classic delayed, visually-guided saccade task with a small spot as the saccade target. In fact, we used this condition as our “spot” condition in Results. Also note that because high spatial frequencies are also associated with reduced visibility (due to the contrast sensitivity function) as well as weakened SC visual responses^17^, the 11 cpd condition from the Saccades-to-Spatial-Frequency paradigm above was also visually quite similar to a classic spot paradigm; it was thus more like a broad-band stimulus rather than a narrow-band one. This explains some of the neuron preference histograms of early stimulus-evoked visual bursts that are shown in Results (e.g. Fig. 3D, E, visual epoch).

We also ran a Saccades-to-Orientation version of the same task. In this case, both the spatial frequency (1 cpd) and contrast (100%) were fixed across trials. However, the grating could have different orientations. We tested 4 orientations across trials, with the following convention of defining orientation: horizontal was defined as 0 deg, and 45, 90, and 135 deg, respectively, were counterclockwise rotations from horizontal.

For Saccades-to-Objects, we were interested in whether our recent observation of visual object detection in the SC^30^ extended to the saccade motor burst epoch. We had images of objects as the eccentric images. Each image was now in a square shape rather than a circular aperture as in the task variants above, and we actually analyzed the motor burst data from the same experiments conducted for our recent work^30^. In that work, we only analyzed the visual burst data at image onset; here, we analyzed the saccade motor bursts. In each session, we had a total of seven natural images, one from each of seven object categories: human face, human hand, monkey neutral face, monkey aggressive face, snake, fruit, and artificial object. We also had another seven images with a grid of horizontal and vertical lines in front of them (as if the object was behind a wire mesh); another seven images with the phase information scrambled but the spatial frequency and luminance content unaltered; and another seven images with grid scrambling, or a random reshuffling of the grid locations from the image with the objects behind a grid of horizontal and vertical lines^30^. The procedures for obtaining all 28 images were described recently^30^, and we used existing toolboxes^61^ to equalize the images in terms of luminance and spatial frequency content^30^. Thus, in total, the monkey made saccades to one of 28 different images in this version of the paradigm. Across sessions, we generated new images that were not used in the previous sessions.

In all of the above task variants, there was a forced delay between target onset and saccade onset. However, in natural behavior, image onsets might “reflexively capture” saccades immediately, meaning that the visual and motor SC bursts occur much closer to each other in time. Therefore, to test the generalizability of our results to immediate, visually-guided saccade situations, we designed a second Saccades-to-X paradigm, but now without a forced delay. The monkey fixated a central fixation spot. After 600-1500 ms, the fixation spot was removed, and an eccentric gray target appeared simultaneously. We used a new feature dimension in this variant of the task, in order to demonstrate the robustness of the phenomenon of sensory tuning in SC neuronal movement bursts. The feature that we used this time was luminance contrast polarity: targets darker than the background were defined as negative polarity stimuli, and targets brighter than the background were defined as positive polarity stimuli^19,62^. The targets consisted of discs of 0.51 deg radius, and they had one of three absolute Weber contrasts (10%, 50%, and 100%). Thus, in this Saccades-to-Luminance-Polarity task, we had six conditions: two luminance polarities (dark versus bright), and three Weber contrasts per polarity. Also note that in this task, we did not provide a central marker spot on the discs as in the above tasks, and this was because the discs were relatively small already; we chose such small discs to demonstrate that the results from the above experiments were not specific to only larger images. Visual bursts from this task were analyzed in our recent study^19^, but saccade-related motor bursts were not inspected.

We collected 50 repetitions per condition per session from the Saccades-to-Spatial-Frequency, Saccades-to-Contrast, Saccades-to-Orientation, and Saccades-to-Luminance-Polarity tasks, and we collected 30 repetitions per condition per session from the Saccades-to-Objects task (because of the increased number of conditions in this task). We typically collected Saccades-to-Spatial-Frequency, Saccades-to-Contrast, and Saccades-to-Orientation within the same session. However, Saccades-to-Objects required dedicated sessions due to the larger numbers of trials that were needed. The Saccades-to-Luminance-Polarity tasks were run, largely, in separate sessions as parts of other experiments, like those described in our recent work^19^. Finally, for the V1 recordings, we ran the Saccades-to-Spatial-Frequency, Saccades-to-Contrast, and Saccades-to-Orientation tasks, and in simultaneous recording sessions of both the SC and V1; the gratings were placed such that overlapping visual response fields (RF’s) of the recorded neurons were visually-stimulated.

In all cases, we placed the saccade target at our estimated best RF and/or mRF hotspot location of the recorded neurons, and we maintained its position throughout a block. This meant that we ran RF mapping tasks before the main paradigms. These tasks were often the classic delayed, visually-guided saccade task or a fixation variant of it, in which no saccade at the end of the trial was required^30^. We sometimes also ran memory-guided saccades, but for other purposes^12^. Our choice of grating size was to approximately fill classic SC visual RF’s at the eccentricities that we tested. For simultaneous SC and V1 recordings, this meant that the grating was larger than V1 RF’s, since we observed that V1 visual RF’s were smaller than in the SC. However, the gratings still very robustly activated V1 neurons, as seen in Fig. 6 in Results.

### Animal eye movement data analysis

We detected saccades in all trials using our established methods^63,64^.

To ensure that we were comparing neuronal activity with similar saccadic execution across all different image types within a given paradigm (e.g. Saccades-to-Contrast), we first ensured that saccade vectors were matched across the image types of the block before proceeding with any neuronal comparisons. This is because SC movement-related RF’s are organized topographically^3,65^; therefore, if one image systematically elicited slightly different saccade vectors from another image, then two different parts of a given neuron’s movement-related RF would be activated by the two images, rendering any differences in motor burst strengths trivially explained by a difference in the saccade vectors. As a result, we always first ensured that we were comparing motor bursts for vector-matched saccades across all image manipulations within a given task block. For example, for Saccades-to-Contrast, we collected all saccades for each image contrast. We then binned all vector endpoints of the saccades into a binning grid with 0.5 deg resolution (in each of the horizontal and vertical directions). We only included trials into any subsequent analyses if a given vector bin had saccades from all image types in the blocks (i.e. all contrasts in the example of Saccades-to-Contrast). In our example of the Saccades-to-Contrast task, if a binning grid location had only saccade vectors from low contrast images but not high contrast images, then this would mean that the saccade vectors for low and high contrasts were slightly different from each other. These saccades were, therefore, excluded from any further analyses. Because we provided a central marker on most images to guide the saccades during the experiments, we still ended up with sufficient trial repetitions for the analyses after the vector matching procedures, as evidenced by the individual example trial rasters shown in Figs. 1, 4 and Supp. Figs. 3, 6.

To further rule out subtle systematic differences between saccades to one image type versus another as the trivial explanation of our results, after matching the vectors of the saccades per the above procedure, we proceeded to analyze the movements’ metrics and kinematics across image conditions. For metrics, we calculated the direction error and the amplitude error of each saccade. Direction error was defined as the angular difference between the vector of the executed saccade and the vector of the image center relative to fixation; amplitude error was defined as the difference in radial amplitude between the vector of the executed saccade and the radial eccentricity of the image center (the saccade target). For kinematics, we calculated saccadic peak velocity. For a given saccade amplitude (as in our task design), the peak velocity should be relatively constant because of the well-known saccadic main sequence relationship^66^ (the monkeys were equally rewarded across trials, so other variables that influence saccade speed, like reward, were equalized). Thus, if the peak velocity is similar for saccades to different image types and the SC motor bursts are very different, then this represents a clear dissociation between SC neuronal activity at the time of saccades from the control of saccade execution, as we and others also observed earlier^12,13^.

For Saccades-to-Objects, analyses of saccade metrics and kinematics (e.g. Fig. 4D, E and Supp. Fig. 8) convinced us that the eye movement properties were already well matched across conditions. Therefore, we analyzed all trials in this task, skipping post-hoc vector matching filters. This was important because there was a larger number of conditions to run in this task, and because vector matching might have caused severe biases in which images were included or removed as opposed to others in a given analysis.

As we describe in more detail below, we typically grouped trials according to the SC motor burst strength. For example, if a neuron had the strongest motor burst for a high contrast image as the saccade target, we defined this image as the “most preferred image” for the “motor burst”. Similarly, if the same neuron had the weakest motor burst for a low contrast image, then this image was the “least preferred image” for the “motor burst”. With such classification, we could compare saccade metric and kinematic properties for the most and least preferred trials. Therefore, we also analyzed the saccades of such trials. We always showed full distributions of our data points, and we also included descriptive statistics (e.g. mean or median values). For kinematic comparisons between most and least preferred images, we additionally calculated a kinematic modulation index. This index was defined as the peak speed of the eye for saccades to the most preferred image minus peak speed for saccades to the least preferred image, divided by the sum of peak speeds. Thus, if the kinematic modulation index was zero, it meant that saccade peak speed was the same for trials with the most and least preferred images. For each neuron, we had a kinematic modulation index from its sessions’ saccades, which we compared to a neuronal modulation index described below. Note that we often recorded multiple neurons simultaneously.

However, since different neurons could have different most and least preferred images, the saccades used for computing kinematic modulation indices (or for other plots of saccadic behavior) were not necessarily the very same saccades for multiple simultaneously recorded neurons.

### Neuronal data analysis

We sorted individual neurons offline using the Kilosort Toolbox^67^, followed by manual curation using the phy software. We then proceeded to analyze spike times and firing rates in the different conditions. To obtain firing rates, we convolved spike times with Gaussian kernels of σ 10 ms.

Our primary goal was to analyze saccade-related “motor” bursts in the SC. To do so, we defined a motor burst epoch as the time interval between −50 ms and 25 ms from saccade onset, in which we measured average firing rates. For comparison, we also analyzed stimulus-evoked visual bursts occurring immediately after image onset. For those, we defined a visual burst epoch as the time interval between 50 ms and 150 ms after image onset during gaze fixation, and we measured average firing rate in this interval.

For classifying SC neurons into different functional cell types (e.g. visual, visual-delay, visual-motor, or motor), we also had additional measurement intervals. The baseline interval was defined as −100 to 0 ms relative to image onset at trial beginning, and the delay-period interval was 400-500 ms after image onset. Within each task variant that we analyzed (e.g. Saccades-to-Contrast), we classified each neuron as being predominantly visual (exhibiting only stimulus-evoked visual bursts), predominantly motor (exhibiting only saccade-related bursts), visual-motor, or visual-delay (exhibiting visual and saccade-prelude activity but no significant motor activity); see Fig. 5A for examples. To do so, we used the measured firing rates in the four measurement epochs described above (baseline, visual epoch, delay epoch, and motor burst epoch) and computed a non-parametric ANOVA (Kruskal-Wallis). We then determined neuronal class by post-hoc tests at the p<0.05 level. Very few well-isolated neurons were not classified into any of the above categories, whether due to low activity levels or other reasons causing the statistical tests to fail. In our population analyses pooling all neuron types, we also included these minority unclassified neurons because inspecting them revealed the same patterns as those of the well-categorized neurons. Also note that we classified neurons separately in each task variant because our primary goal was to ask whether sensory-tuning in SC neuronal movement commands was robust even in neurons with relatively stronger movement-related rather than visual-related activity (e.g. Fig. 5) within any given task. The question of whether SC mRF’s themselves are different for different image types is orthogonal to this investigation and requires dedicated mRF mapping sessions with multiple image types (a technically-challenging endeavor due to the numbers of trials required). Moreover, there could be task-related variability in firing rates (e.g. differences in delay-period activity) across different stimulus types. Most importantly, our neuron classification was highly robust across the different tasks, as evidenced by the similarity of our visual-motor indices (described below) across tasks (e.g. Supp. Fig. 9A).

For analyzing SC saccade-related “motor” bursts, in each task variant (e.g. Saccades-to-Contrast), we defined (for each neuron) the image associated with the strongest saccade-related motor burst as the “most preferred” image. We also defined the image associated with the weakest saccade-related motor burst as the “least preferred” image. This was done after the vector-matching procedures described above. Because different neurons had different preferred and non-preferred images (see Results), this classification allowed us to obtain population-level effect sizes across neurons. To do so, we normalized each neuron’s firing rate and then averaged across neurons. Normalization was done as follows. In each neuron, we found the peak firing rate occurring either after stimulus onset or around saccade triggering. We then used the larger of the two peaks and subtracted the neuron’s baseline activity from it. This constituted our normalization constant. For any firing rate that we wanted to normalize in the neuron’s data, we subtracted the neuron’s baseline activity from it and then divided by the baseline-subtracted maximal response of the neuron (i.e. by our normalization constant). After obtaining the population saccade-related motor burst strengths for the most and least preferred images, we plotted the differences between them as well (e.g. Fig. 2A, gray).

We also calculated neuronal modulation indices, similar to what we did with the kinematic modulation indices described above. For each neuron, we plotted the average firing rate curve of the neuron around saccade onset. We then measured the maximum of this curve in the interval from −50 to 25 ms relative to saccade onset, and we did this for either the most or least preferred feature. The neuronal modulation index was defined as the burst strength of the neuron for the most preferred feature minus the burst strength for the least preferred feature divided by the sum of the two. We then plotted the distributions of neuronal modulation indices across neurons in our different tasks. To compare neuronal modulation indices to kinematic modulation indices, we plotted the two indices against each other for each task, and we calculated correlation coefficients. This allowed us to assess whether a large change in burst firing rate in a given neuron (e.g. Fig. 2A) was associated with an equally large change in saccade kinematics or not.

Because we found that the most and least preferred images could be different within a single neuron between the early stimulus-evoked visual burst epoch and the saccade-related motor burst epoch (e.g. Supp. Fig. 6A, B), we also repeated the above procedure for the early “visual” epoch of the trials (immediately after image onset during fixation), but after first identifying the most and least preferred images of each neuron in this visual epoch. The normalization of firing rates was not re-done since the above normalization was applied to entire trials and not just the motor burst epochs.

We also performed comparisons on raw firing rates, either by plotting the raw measurements directly (e.g. Fig. 2B), or by using receiver-operating-characteristic (ROC) analyses in a manner similar to other studies^20^. We calculated the area under the ROC curve (AUC) as a function of time in either saccade or stimulus-evoked visual epochs. For each trial of the “most preferred” image, we measured instantaneous firing rate at a given point (e.g. near saccade onset), and we did the same for the “least preferred” image. We then calculated the AUC across the distribution of trials at that time point. We repeated this procedure as a function of time, and this gave us time courses of AUC changes relative to either saccade onset or stimulus onset. This procedure allowed us to demonstrate differences in saccade-related bursts despite matched saccade vectors, and it is similar to analyses performed for pre-saccadic elevations in visual cortical neurons of area V4^20^. In some analyses, we also performed AUC analyses as a function of identified cell type. Here, we used the classification of neurons described above and performed the analysis only on neurons within a given functional cell class. Our AUC calculations were similar to those we used recently^30^.

For Saccades-to-Objects, we were particularly struck by the preference of saccade-related motor bursts to real object images as opposed to scrambles (e.g. Fig. 4F). Therefore, we checked whether neurons had significant peri-movement AUC elevations when comparing object images to scrambled images. Thus, we grouped all seven image categories into one. This resulted in four groups: real objects, grid-covered real objects, phase-scrambled objects, and grid-scrambled objects. We then checked for significant peri-movement AUC values when comparing real objects to either phase or grid-scrambled categories (or both). We assessed significance, similarly to how we did it recently for SC visual responses to objects^30^. Specifically, we calculated bootstrapped confidence intervals for AUC measures; neurons that had AUC values significantly different from 0.5 at the p<0.05 anywhere from times −100 to 100 ms relative to saccade onset were deemed to be significant.

For local field potential (LFP) analyses, we obtained raw wide-band signals from each electrode contact. We then applied zero-lag filtering procedures as described previously^68^. Briefly, we used notch filtering to remove the line noise frequency (50 Hz) and its next two harmonics (100 and 150 Hz), and we also kept signals <300 Hz as the LFP band. To classify whether the channel from which we collected LFP’s was from the more visual (superficial) or more motor (deep) SC layers, we classified each electrode channel’s multi-unit activity (MUA) as being predominantly visual or predominantly motor using a visual-motor index (VMI)^51^. Specifically, for each channel, we filtered the wide-band signal using a fourth-order Butterworth band-pass filter (750 to 5000 Hz), and we then rectified the signal before passing it through a second low-pass filter (fourth-order Butterworth) with 500 Hz frequency cutoff. For each condition, we plotted stimulus- and saccade-aligned MUA responses after subtracting the baseline MUA level (defined as the average MUA in the final 200 ms before image onset); superficial channels had stronger visual than motor responses, whereas deeper channels had stronger motor than visual response^51^. To quantify this, we measured a motor MUA value and a visual MUA value. These were defined as the average baseline-subtracted MUA in the interval −25 to 25 ms from saccade onset (for the motor MUA measurement) or 30 to 200 ms after image onset (for the visual MUA measurement). The VMI was defined as the motor MUA measurement minus the visual MUA measurement divided by the sum of the two. VMI’s larger than zero were more motor than visual (e.g. Supp. Fig. 9A).

For state-space analyses, we performed a pseudo-population analysis^26,69^. For each task, the instantaneous firing rate of all neurons that we recorded from was a point in an N-dimensional space of the activity of the population of N neurons. As all neurons’ firing rates changed across time (e.g. after stimulus onset or peri-saccadically), the population activity representation moved in this N-dimensional space. We, thus, assumed stability across sessions of SC activity since not all neurons in our population were recorded simultaneously^26^. Since population activity likely occupied a much lower dimension than the number of neurons, we performed principal components analysis (PCA) and plotted the population trajectory within the first 3 PCA dimensions. These typically accounted for the majority of the variance of population firing rates (e.g. 70-89%). We used such state-space analysis to first compare visual and motor burst population trajectories and then to check for tuning in the motor bursts. In both cases, we normalized each neuron’s firing rate before performing PCA, using the same normalization procedure described above.

To compare visual and motor burst population trajectories in PCA space, we concatenated each neuron’s activity in a visual interval (from 0 to 200 ms relative to stimulus onset) with activity in a saccade epoch (from −100 to 50 ms relative to saccade onset). Then, we projected the population activity on 3-dimensional PCA space. This allowed us to assess whether visual and motor SC activity occupied similar or different subspaces (e.g. Fig. 3G).

To check for sensory tuning in the motor bursts themselves, we focused on the peri-saccadic interval only, and we projected SC population activity in this interval, using PCA, for different images as the saccade targets. If there was sensory tuning in the SC population motor bursts, then the population peri-saccadic state-space trajectories should differ as a function of which image was the saccade target (despite the vector- and kinematically-matched saccades) (e.g. Fig. 3H). To quantitatively confirm such difference, we then picked a reference peri-saccadic trajectory from one of the image features of the experiment (e.g. spot in the Saccades-to-Contrast experiment), and we calculated the Euclidean distance of each other image feature’s population activity trajectory from this reference trajectory. We did this for times around saccade onset. Moreover, we calculated Euclidean distances from the entire high-dimensional population space, and not just from the 3-dimensional PCA sub-projection of only 3 principal components. For checking Euclidean distances against a null distance distribution, we performed 1000 permutations in which we randomly picked a reference and a condition trajectory, and we then calculated theEuclidean distance between the two.

Analyses of V1 visual responses were similar to those of SC visual responses, except that our measurement interval was 30 to 150 ms after stimulus onset, since we observed that V1 neurons had slightly earlier visual response latencies than SC neurons.

### Human psychophysical experiments

Our SC results suggest that SC motor bursts contain information about the saccade target’s visual features. Since such bursts are relayed to the cortex virtually unchanged^31^, this suggests that peri-saccadic corollary discharge^32,34,70^ from the SC may relay not only the movement vector information, as previously suggested, but also information about the visual appearance of the saccade target. If so, then perceptual phenomena that are believed to depend on peri-saccadic corollary discharge, such as peri-saccadic perceptual suppression^33^ and peri-saccadic perceptual mislocation^34^, may be modulated by the visual appearance of the saccade target. We, therefore, ran these two paradigms (suppression and mislocalization) on 8 human subjects each (7 naïve and one author).

The experiments were approved by ethical committees at the Medical Faculty of the University of Tübingen, and the subjects provided informed written consent. They were also financially compensated for their time, and the experiments were in line with the Declaration of Helsinki.

In the peri-saccadic suppression experiment, we tested 8 subjects (5 female), aged 23-27 years. In the peri-saccadic mislocalization experiment, we tested 8 subjects (4 female), aged 23-29 years. Four of these subjects had also performed the peri-saccadic suppression experiment in previous sessions.

The laboratory setup was the same as used in our recent studies^44,71^, and we largely used similar procedures. Briefly, the subjects sat in a dark room 57 cm from a calibrated and linearized CRT display with 41 pixels/deg resolution. We tracked their eye movements at 1 KHz using a video-based eye tracker (EyeLink1000, SR Research), and we stabilized their head position using a custom-built apparatus described earlier^72^. Stimuli were presented on a gray background of 20.01 cd/m^2^ luminance, and fixation spots (7 by 7 min arc) were white (with 49.25 cd/m^2^ luminance).

In the peri-saccadic suppression experiment, the subjects fixated a white spot placed ∼8.5 deg either to the right or left of display center. After 500 ms, a low (0.5 cpd) or high (5 cpd) spatial frequency Gabor grating was presented at screen center. The grating had a gaussian smoothing window with σ 0.49 deg, and it had high contrast (100%); its phase was random. The subjects were still required to maintain fixation for another 1000-2000 ms (because we wanted to avoid any sensory transients associated with grating onset). When the fixation spot was removed, the subjects made a saccade towards the center of the grating, which had a central marker in its middle like in the neurophysiology experiments (to minimize inter-trial saccade variance). Once we detected a saccade onset, as per the procedures we described recently^44,71^, we presented a brief (1-frame; 85 Hz refresh rate in the display) flash of a square of size 45 by 45 min arc and variable luminance from trial to trial. The flash time was either immediate upon online saccade detection, or at two later times (+11 or +35 ms). In post-hoc analyses, we recomputed the flash times to always report them relative to actual saccade onset (histograms shown in Supp. Fig. 10B). The flash could appear at one of four cardinal locations relative to the center of the display (and at an eccentricity halfway from the initial fixation position). The subjects’ task was to report which of the four locations had a brief flash via a button box. Across trials, we varied the flash luminance to obtain full psychometric functions of flash detection sensitivity. The procedure was similar to that we described recently^44^. Briefly, we performed a staircase procedure in the first session, to obtain an initial estimate of perceptual threshold in a subject, and in subsequent sessions, we also added additional samples of flash contrast to sample more locations on the psychometric curves. Importantly, we wanted to avoid having references frames (e.g. the edges of the display monitor) strongly influence performance. Therefore, across trials, we randomly shifted all stimulus positions in a trial by a random number from the range of +/-1.2 deg in either horizontal or vertical direction (at pixel resolution; 1/41 deg). This imposed position variance minimized the possibility of subjects remembering the absolute position of flashes relative to the display monitor edges. We collected approximately 450-500 trials per spatial frequency in each subject.

The peri-saccadic mislocalization experiment was very similar except that the brief flash was always a bright white square (high contrast). The times of the flash were also adjusted slightly to be: immediately upon saccade detection or at +40 or +80 ms from online saccade detection. Moreover, the task of the subjects was to point (with a computer mouse) to the perceived location of the flash. Specifically, after the subjects fixated the central grating for 500 ms, we removed all stimuli from the display, and displayed a mouse cursor (at display center). The subjects clicked on the perceived flash location, and they were free to move their eyes until they responded (they typically made a second saccade towards the perceived flash position). We picked four possible flash locations, each at a radius of ∼7.3 deg from the saccade target center location. One flash was directly opposite the saccade target direction (i.e. on the horizontal axis), and the other three flashes were along the saccade direction: one on the horizontal axis and the other two at the +/-45 deg diagonals. Even though the flash opposite the saccade vector was expected to have the least mislocalization^73^ (which we confirmed), we included it so that subjects could not expect always seeing flashes ahead of the saccade target. In the analyses, we only analyzed the flash locations ahead of the saccade target to check whether mislocalization was different for low or high spatial frequency saccade targets (e.g. Supp. Fig. 10G-J). We collected 320 trials per spatial frequency in each subject.

Note that in both experiments, the low and high spatial frequencies chosen as the saccade targets were differentially represented by the SC motor bursts in our neurophysiological results (Fig. 3D, Saccade epoch and Fig. 3H, I). Therefore, we hypothesized that the peri-saccadic perceptual effects might be different for low and high spatial frequencies.

### Human psychophysical experiments’ data analysis

We detected saccades and microsaccades as described above. Before analyzing any perceptual reports, we filtered the eye movement data to ensure matched saccade execution across low and high spatial frequency saccade targets. First, we defined a maximal radius (2 deg) for initial eye position at saccade triggering, and made sure that trials with both high and low spatial frequency saccade targets had the same initial starting positions. Similarly, we defined a maximal radius for final eye position at the end of the saccade, to make sure we had matched saccade vectors. Then, for each saccade direction, we ensured that the amplitudes and peak velocities of the saccades were the same regardless of whether the saccade target had a high or low spatial frequency (e.g. Supp. Fig. 10C, F). We also excluded trials with saccadic reaction times <100 ms or >500 ms. Finally, we recomputed all flash times relative to the actual saccade onset times after we properly detected the movements.

To obtain psychometric curves (in the peri-saccadic suppression experiment), we used the Psignifit 4 toolbox^74^ for each subject individually, using a cumulative gaussian function (and 0 lapse rate). We then averaged across the subjects’ individual psychometric curves to obtain a population measure. We did this for each time bin of flash times relative to saccade onset (with the time bins defined in Supp. Fig. 10B as t1, t2, and t3, respectively).

Statistically, we performed bootstrapping to check whether perceptual suppression depended on the saccade target image type. To do so, we created random permutations (10000) in which we randomly picked trials for the two conditions (low or high spatial frequency saccade target). When doing so, we kept the same numbers of observations per subject per condition, in order to maintain the sampling numbers consistent with our actual experimental data. We then performed psychometric curve fits from such permuted data and measured the threshold at the 62.5% correct rate. Across the 10000 permutations, we plotted the distributions of thresholds between the two sets of permuted conditions. If our actual measured threshold difference between low and high spatial frequency saccade targets was deviated by more than 95% of the threshold differences of the random permutations, then we deemed our measured threshold difference in the real experiment to be significant. We found that there was a significant difference between the low and high spatial frequency targets at the first time bin (t1) in which flashes happened relative to saccade onset (Supp. Fig. 10E). No significant differences were observed at longer times from saccade onset.

For peri-saccadic mislocalization, we first confirmed that we had matched saccade vectors like in the peri-saccadic suppression experiment. We then confirmed that saccade peak velocities were similar for low and high spatial frequency saccade targets (e.g. Supp. Fig. 10F). We then collected each subject’s click locations at different flash times. To summarize these click locations, we picked the median click location for each subject and each condition (low or high spatial frequency saccade target). We found that mislocalization was extreme at the shortest flash time, causing ceiling effects (clicks very near the saccade target). Therefore, we analyzed click locations at flash time around 70 ms from saccade onset, and we compared them to clicks for a longer flash time (around 110 ms). To summarize the mislocalization effect, we measured the Euclidean distance from the actual flash location to the median click location in each subject. Thus, larger mislocalization was associated with a larger Euclidean distance. We then plotted the mean and s.e.m. of the distance across subjects, and we showed descriptive statistics and underlying individual subject results (e.g. Supp. Fig. 10I, J).

